# The evolution of an individual-like dispersive stage in colonial siphonophores

**DOI:** 10.1101/2024.07.15.603641

**Authors:** Maciej K. Mańko, Catriona Munro, Lucas Leclère

## Abstract

Evolutionary transitions between individual and colonial organisms remain enigmatic. Siphonophores, abundant pelagic cnidarians, exhibit a complex colony structure composed of repeated individual (zooid) clusters called cormidia. Many siphonophores release their posterior-most cormidia as independent fragments known as eudoxids, ensuring sexual reproduction. However, the mechanisms of eudoxid production and its evolutionary origins are unknown. Using live imaging, immunohistochemistry and pharmacological inhibition we provide a mechanistic understanding of eudoxid formation. We demonstrate that eudoxid release is controlled by a dedicated muscle and involves tissue remodeling, leading to the formation of an integrated dispersive unit with specific behaviors and different buoyancy. We show that eudoxids and parental colonies often have different spatial or temporal distributions, suggesting niche partitioning. We infer that eudoxids evolved once through the concomitant evolution of multiple cormidium subparts. This study reveals how the acquisition of an individual-like dispersal stage, through muscle evolution and colony modification, contributed to the ecological success of a primary carnivore in marine planktonic ecosystems.

## Introduction

The hierarchical integration of lower-level entities into higher-level units of selection has led to some of the most remarkable cases of biological evolution, from multicellular aggregations of cells, to colonial or social aggregations of individuals^1–3^. As the integration progresses, lower-level units may specialize, and their individuality may become subordinated^4^. In clonal colonial animals, such as hydrozoans (Cnidaria), this evolutionary transition in individuality has been likely achieved by retention of asexually budded individuals, driving subsequent division of labor and polymorphism of colony-building modules^5^, followed by an evolutionary complexity drain^6^. This functional diversification of modules, integrated to collectively function a single organism, blurs the distinction between the individual and the colony.

Clonal colonial organization, in which genetically identical individuals (called zooids, ramets, etc.) remain physiologically integrated, offers multiple ecological advantages, derived from their modularity and unconstrained growth^7^. Examples of colonial taxa are particularly abundant among sessile marine fauna^6^ (*e.g*., bryozoans and corals), whose immobility limits dispersal capacity and thus increases risks of intraspecific competition and inbreeding^8,9^. To overcome these challenges, colonial species have evolved complex life cycles with a dispersive stage either before fertilization (a reproductive individual released from the colony) or after (planktonic larvae), enhancing their ability to disperse over longer distances despite their sedentary lifestyle. However, not all colonial taxa are sessile, with planktonic colonies found in two animal phyla: Chordata (salps, doliolids and pyrosomes) and Cnidaria (porpitiids and siphonophores). Unlike their benthic counterparts, planktonic colonies are characterized by directional growth with limited plasticity or branching^4^, and are motile^10^, thus have broader distributional ranges^11^. Despite their greater dispersal capacity, planktonic colonial species still display complex life cycles, the ecological and evolutionary significance of which remain to be addressed.

Siphonophores, a clade of entirely marine hydrozoan cnidarians, have unequivocally the most complex colony architecture and the highest degree of zooid functional specialization among animals^12^. The majority of siphonophore colonies are streamlined, with asexual budding of zooids, homologous to free-living polyps and medusae, restricted to either one or two localized growth zones^12,13^. Non-locomotory zooids are budded off from a single growth zone, giving rise to iterative clusters of polymorphic zooids, called cormidia, dispersed in an ontogenetic sequence along an elongated stem^4,13^. Despite limited data on siphonophore reproduction and life cycles^14^, siphonophores are known to decouple sexual reproduction from parental colonies, by detaching gonad-bearing reproductive individuals, gonophores, either individually or in clusters^15^.

Release of reproductive individuals is well documented across hydrozoans, where canonical life cycle involves detachment of sexually reproducing individual, medusa, from a benthic polyp. Siphonophore gonophores, being motile but not able to feed, fall centrally within hydrozoan continuum of medusa truncation, which ranges from fully independent medusa (moving and feeding), to sporosacs, which typically remain attached to polyps^16^. Some members of the most-speciose clade of siphonophores, Calycophorae, have taken the reproduction related specialization one step further. Instead of releasing just the reproductive individuals (gonophores), their colonies undergo evolutionarily unique, sequential fragmentation, detaching entire cormidia – called eudoxids after release – which are comprised of the gonophore and a number of other functionally specialized zooids. This renders organismal nature of eudoxid puzzling^17^. If assessed by their multi-zooid composition, they could be considered either a colony, albeit incapable of budding (other than of gonophores) and originating from another colony^18^, or as individuals^19^. However, this intriguing interpretation would require evidence of the sufficient physiological and functional integration of eudoxid components, enabling it to function as an autonomous and integrated unit^3,20,21^, a requirement that is largely unaddressed.

Eudoxid exhibit distinct morphology compared to the cormidia still attached to the parental colony. This has led earlier authors to describe eudoxids and their parental colonies as separate species^22^, much like the case of medusae and their parental polyps in other cnidarians^23^. The realization that the eudoxid represents a life cycle stage rather than a different species came already in 1853^24,25^. Nevertheless, new “eudoxid species’’ continued to be described, under the genus *Eudoxia*^26^, as the morphological distinctiveness of eudoxids and cormidia still attached to parental colonies hampered unification of these two stages under a single name. Furthermore, the extreme similarity of eudoxids, but not of parental colonies, in congeneric species, made such definitive linking possible only if the molecular tools were applied^27^.

Despite over two centuries of siphonophore research^13^, the fragility of siphonophore colonies and the inability to culture them^14^ precluded a thorough characterizing of eudoxid production. The sole attempt at describing this process, conducted in 1871^21^, provided insightful suggestions on stem narrowing at week points, laying the foundation for our work. Here, we provide a mechanistic understanding of eudoxid production and uncover its ecological significance and evolutionary history. We describe stages of eudoxid development and release using *in vivo* imaging, followed by documenting cellular dynamics through immunohistochemistry, electron microscopy and pharmacological inhibition experiments. We also assess ecological significance of eudoxid acquisition by reviewing the entire siphonophore literature, describe a new eudoxid behavior, reconstruct the evolution of colony organization and eudoxid morphology based on an improved siphonophore phylogeny.

## Results

### Stages of cormidium development

We characterized eudoxid formation in *Chelophyes appendiculata*, a highly abundant, epipelagic siphonophore species, known to release eudoxids^28^, whose ease of collection permitted extensive observations. The formation comprises three spatially and temporarily separated processes: cormidium development, cormidium release and eudoxid maturation, of which the latter occurs only in released cormidia.

Four distinct stages of cormidium development can be distinguished based on differences in size and morphology of bract and gonophore. Newly budded cormidia appear as an undifferentiated projection from the stem, that develops into a gastrozooid (Fig. 2B, E), thus entering into stage 1 (S1) of cormidium development. As the gastrozooid enlarges and its tentacle develops, the bract buds off from the left-hand side of the gastrozooid’s peduncle (see^29^ for siphonophore axes; Fig. 2A, B, C, E). Actin-rich structures can be seen at the base of the bracteal canal (Fig. 2C), whose presence in other connection sites (*e.g*., stem-nectophore, Supplementary Fig. 1) suggest their role in zooid attachment. The bract, initially a minute, shield-like zooid covering the gastrozooid’s base, progressively extends over the stem migrating towards the stem’s dorsal side (Fig. 2A, B), while the bracteal canal elongates to eventually become positioned perpendicularly to the anterior-posterior axis of the stem. The migration of the bract stops when the central portion of the bracteal canal is opposite to the gastrozooid, coinciding with the appearance of a gonophore bud from the left-hand side of the gastrozooid’s peduncle, marking the end of stage 1 (S1; Fig. 2E, F). As development progresses, the bract enlarges and the gonophore becomes more prominent (Fig. 2F). The bracteal canal, originally a tubular structure embracing the stem, develops an apical projection, marking the transition into stage 2 (S2; Fig. 2F, I). At this stage, the bract resembles a cone wrapped around the stem, with the right side overhanging the left one (Fig. 2F). When the apical projection of the bracteal canal grows to twice the height of its lateral branches, the cormidium enters stage 3 (S3Fig. 2G). At this stage, the gonophore becomes more differentiated and starts pumping actively. The bract’s shape does not visibly differ from the bract at S2. During the transition into stage 4 (S4; Fig. 2H, I), the lateral branches of the bracteal canal retract, while its apical projection enlarges and protrudes deeper into the bract, likely due to the mesoglea build up in the surrounding tissue (Fig. 2H). As a result, only two connections between the bracteal canal and the stem are maintained: the apical connection, joining the bracteal canal’s apical projection with the stem, and the basal one. Simultaneously, the bract shortens and acquires a more triangular shape than in S2 or S3; in addition, the two flaps of the bract retreat revealing bracteal furrow, the groove through which runs the stem (Fig. 2H).

**Fig. 1.**
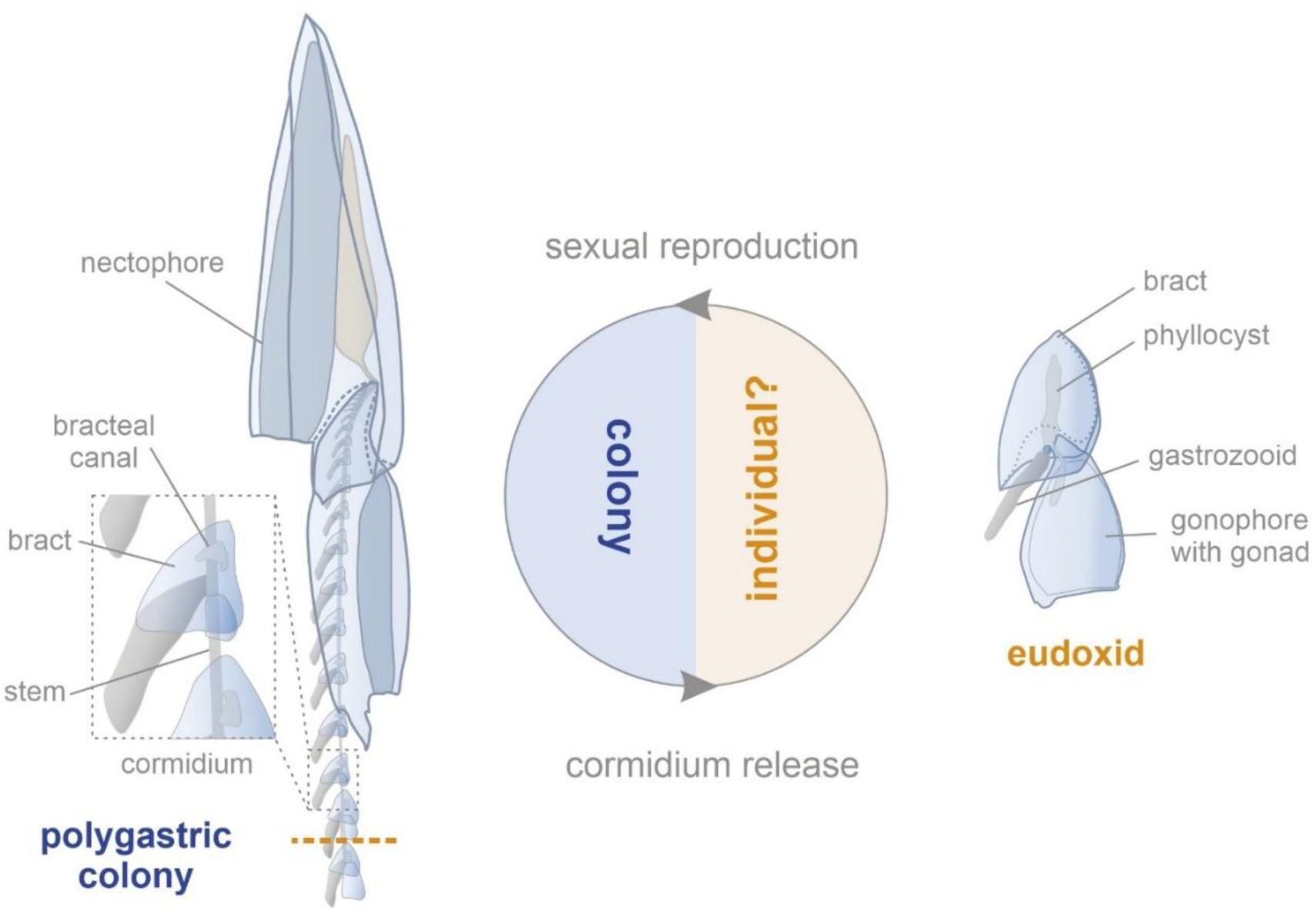
Life cycle and colony-individual transition in calycophoran siphonophore. Life cycle of a calycophoran siphonophore, *Chelophyes appendiculata,* with major processes and morphological features (explained below). Polygastric colony comprises nectophores (swimming bells) an elongated stem (longitudinal stolon connecting colony-members) on which zooids are budded in cormidia (repetitive clusters), each comprising: gastrozooid (feeding zooid), gonophore (gonad bearing zooid) and a bract (gelatinous zooid of unclear homology) with bracteal canal (extension of gastrovascular system). Release of terminal cormidium from polygastric colony gives rise to a dispersive stage, eudoxid, that shows distinct morphology to undetached cormidium in that its bract is of different shape and it contains phyllocyst (swollen branch of bracteal canal). Eudoxid not drawn to scale.

**Fig. 2.**
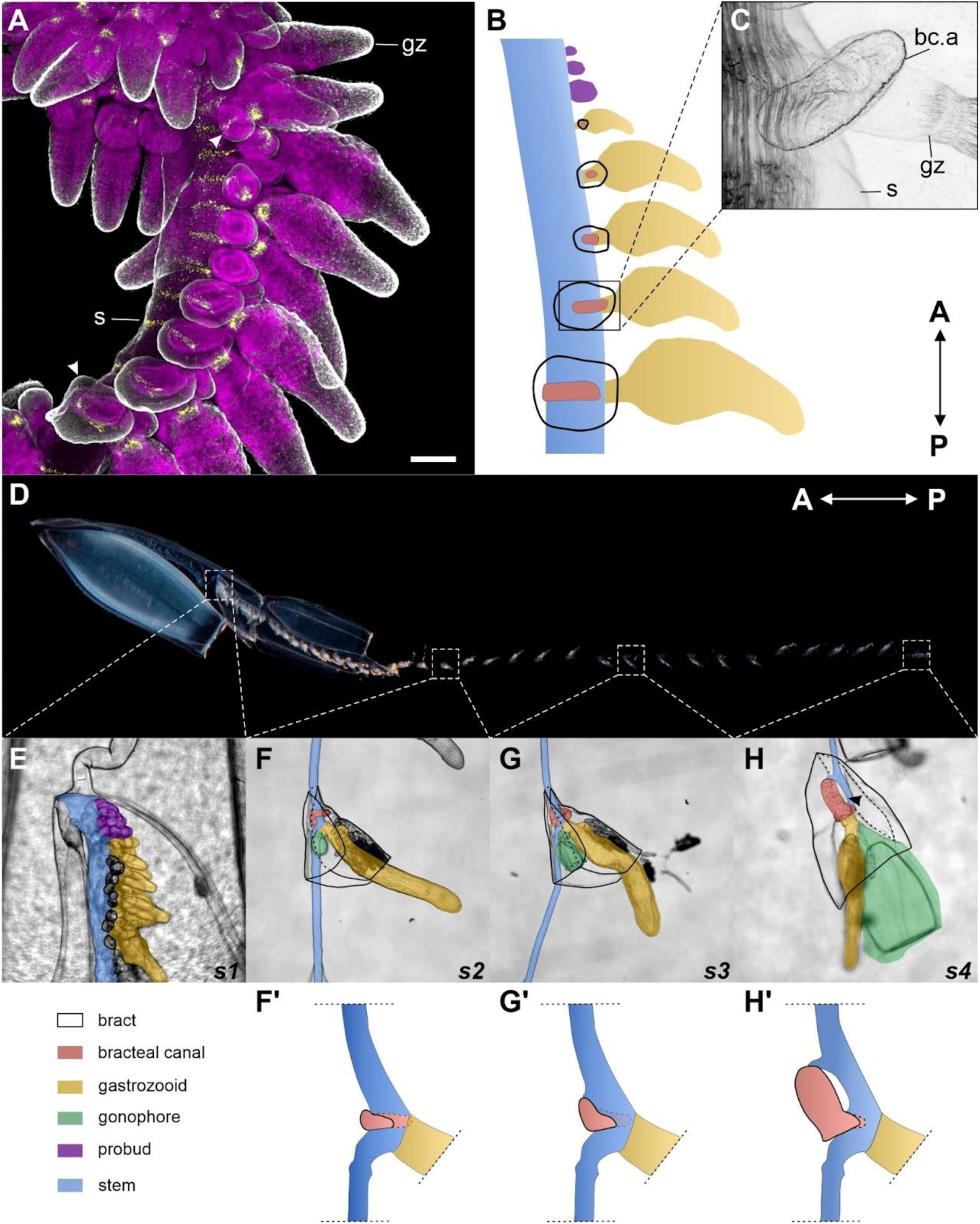
Stages of cormidium development in *Chelophyes appendiculata*. **A.** Immunohistostaining of tyrosinated tubulin (grey), FMFRamide (yellow) and nuclei (magenta) in young *C. appendiculata* stem. Arrowheads point at the youngest (top = anterior) and the oldest (bottom = posterior) bract visible. Scale bar: 100 μm. **B.** Schematic representation of early cormidium differentiation with color-coded zooids. **C.** Actin detection in young stem showing junction of bracteal canal and the gastrovascular system of a gastrozooid. **D.** Overview of *C. appendiculata* colony. **E.–H.** Stages S1–S4 of cormidium development, with color-coded zooids and other components of the colony. Black arrowhead points at the mesoglea build up site between the bracteal canal and the stem. **F’, G’, H’**. Schemes of bracteal canal progression through stages S2 (F), S3 (G) and S4 (H), with the extent of stem and gastrozooid limited by the dashed line; note that the H photo shows terminal S4 cormidium, while the corresponding scheme (H’) documents S4 cormidium morphology in precedent cormidium (*i.e.*, not terminal). Figures are aligned so that the anterior pole of the colony faces up, except for D., where the colony’s anterior pole faces left. Note that figures: A, B, C, E were flipped for presentation purpose, and that in all schemes gastrozooid’s tentacles were omitted for clarity. Labels: bc.a – bracteal canal attachment, gz – gastrozooid, s – stem.

### Disruption of connectivity between cormidia triggers their ordered release

Attempts to characterize cormidium release in actively moving colonies were unsuccessful, as the colonies would randomly fragment upon swimming into the walls of the Petri dish. After immobilizing the nectophores (N_colonies_= 8) or placing the colonies in large containers (N_colonies_= 2), we were able to document that cormidia are released one at a time (20/20 release events), consistently upon reaching S4 of development (14/14; Supplementary Table 1). This process occurred in a fixed order, starting from the posterior end of the colony (20/20).

We also found that cormidium release can be triggered by disrupting the connection between a stem fragment with several cormidia and the remaining colony (Supplementary Table 2). 2-, 3-, and 4-cormidia-long stem fragments (CLSF) at stage S3 (N_CLSFs_ = 1, 8, 7, respectively), dissected off from *C. appendiculata* colonies consistently released all their cormidia (N_released cormidia_ = 56). Interestingly, when a stem from a single colony was cut into two 3-CLSFs (N_colonies_ = 6), the order of cormidia release was maintained within each fragment, but not between the two fragments. The two fragments started releasing their posterior-most fragments nearly simultaneously, suggesting that an inhibitor signal prevents the release of more anterior cormidia.

### Muscle contraction at the detachment ring is involved in cormidium release

To characterize the sequence of events leading to cormidium release, we designed an *in vivo* time-lapse protocol (see Methods), in which 3-CLSFs were immobilized using a glass micro-chamber. Analysis of frame grabs revealed that the release of cormidia is preceded by a constriction of the stem at a specific site, hereafter referred to as the detachment ring. This constriction occurs immediately posterior to the gastrozooid’s peduncle of a second-to-last cormidium of the colony (Fig. 3A, B, Supplementary Movie 1). While the detachment ring remains constricted, the posterior portion of the stem undergoes a series of longitudinal contractions, and the gonophore continues to pump, ultimately leading to the release of the terminal cormidium at the detachment ring. The detachment ring remains constricted even after the release, as indicated by actin staining (Fig. 3D, E), and gets resorbed by the preceding cormidium only later (Fig. 3F). This likely serves to maintain separation of the gastrovascular system from the surrounding environment.

**Fig. 3.**
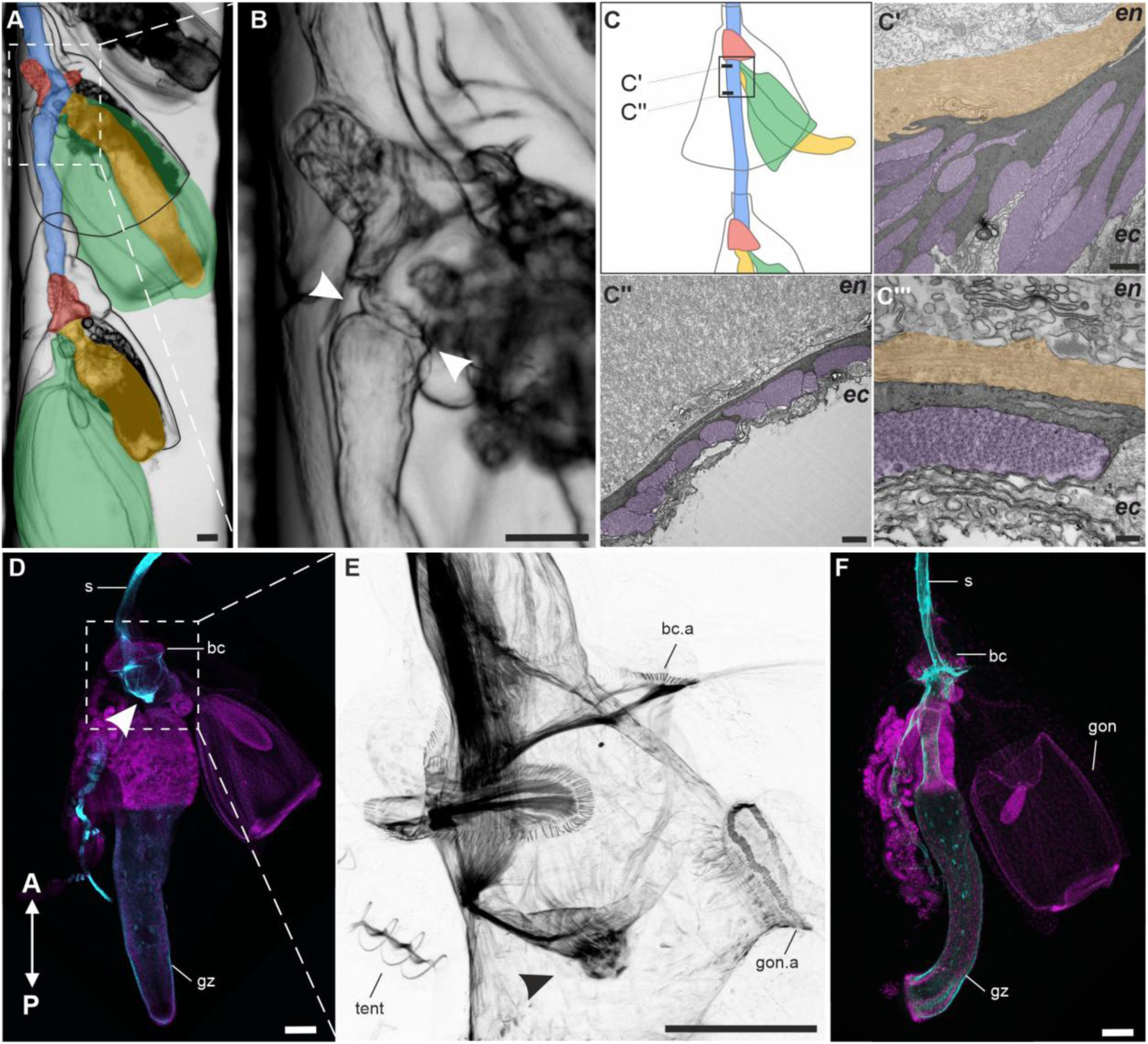
Constriction at the detachment ring is necessary for cormidium release. **A.** A three-cormidia-long stem fragment (from Supplementary Movie 1), with a constricted detachment ring at the second-to-last cormidium. Color coded as in Fig. 1. **B.** Close-up of the detachment ring showed in (A). **C.** Schematics of TEM ultra-thin sections shown in panels C’ and C’’. **C’.** Microanatomy of the detachment ring, with endodermal muscle myofilaments colored in orange and their ectodermal counterparts in purple. **C’’**. Stem microanatomy below the detachment ring, color scheme as in C’. **C’’’.** Close-up of myofilaments across mesoglea within the detachment ring. **D.** Actin (cyan) and nuclei (magenta) detection in the second-to-last cormidium fixed right after the detachment of the posterior-most cormidium. **E.** Close up of the area marked in (D), depicting a constricted detachment ring (arrowhead) and the actin-rich structures associated with attachment sites (Supplementary Fig. 1). **F.** Actin (cyan) and nuclei (magenta) detection in the posterior most cormidium without any visible remnants of the detachment ring. All figures are aligned so that the anterior pole of the colony faces up. Labels: bc – bracteal canal, bc.a – bracteal canal attachment, ec – ectoderm, en – endoderm, gon – gonophore, gon.a – gonophore attachment, gz – gastrozooid, s – stem, tent – tentillum, arrowhead: detachment ring. Scale bars: 100 μm (A, B, D, E, F), 1 μm (C’, C’’), 200 nm (C’’’).

The muscular nature of the detachment ring was corroborated by transmission electron microscopy (Fig. 3C’-C’’’). The detachment ring is made up of two layers of myofilaments, as indicated by the hexagonal lattice of myosin and actin filaments (Fig. 3C’’’): ectodermal and endodermal, separated by an electron dense layer of mesoglea (Fig. 3C’, C’’’). On the ectodermal side, mesoglea branches out, separating individual myofilaments, likely to provide additional support, while no such branching was observed on the endodermal side. Importantly, the two layers of myofilaments are perpendicular to each other (Fig. 3C’’’), with the ectodermal layer running along the stem’s length, while the endodermal layer forms circular bands. In the other parts of the stem, myofilaments were found only in the ectoderm (Fig. 3C’’).

Through pharmacological inhibition experiments on 2- and 3-CLSF we further demonstrate the role of muscle contractions in cormidium release (Supplementary Table 3). Under control conditions, release occurred relatively rapid, with fully formed eudoxids appearing within 22 h in all cases (19/19). Treatment with non-muscle myosin inhibitor slightly impaired stem’s ability to release cormidia, more pronouncedly at higher concentrations (release success of 10/14 and 5/14, for 1 μM and 5 μM Blebbistatin, respectively). In contrast, blocking muscle contractions with an isotonic MgCl_2_ solution consistently impaired the stem’s ability to release cormidia (release success 2/44).

### Release competence correlates with the development of the bracteal canal and detachment ring

Our initial observations indicated that *C. appendiculata* colonies with less developed stems did not produce eudoxids (Supplementary Table 1). To systematically test if this ability correlates with cormidium maturation we dissected 2-, 3- and 4-CLSFs (N_CLSFs_ = 3, 16, 12, respectively) at S2 of development (Supplementary Table 2). Of the 102 S2 cormidia in 31 stem fragments none were released throughout the duration of the experiment, thus indicating that the competence to release arises during the transition from S2 to S3. Interestingly, during this assay bracts detached from all cormidia within the first 24 h of the experiment, suggesting weaker bract-stem attachment at this stage of cormidium development.

We were able to correlate the competence to release eudoxids with the development of the detachment ring and bracteal canal. The detachment ring is already discernible in cormidia within the growth zone (Fig. 4A, A’), with a single detachment ring forming posteriorly to each newly budded gastrozooid. Initially, the detachment ring has a form of a semi-ring, composed of few myofilaments running perpendicularly to the A-P axis of the stem. It extends from the gastrozooids’ base, to roughly the mid-width of the stem on both sides (Fig. 4A, A’). As the cormidium develops, the detachment ring begins to encircle the entire stem (Fig. 4B, B’). Upon entering S3, the detachment ring shifts posteriorly and widens (Fig. 4C, C’), making it distinguishable even without the actin staining (Fig. 4D). Similarly, the bracteal canal changes shape between S2 and S3 (Fig. 2E-H) with a strengthening of its connection to the stem (Fig. 4).

**Fig. 4.**
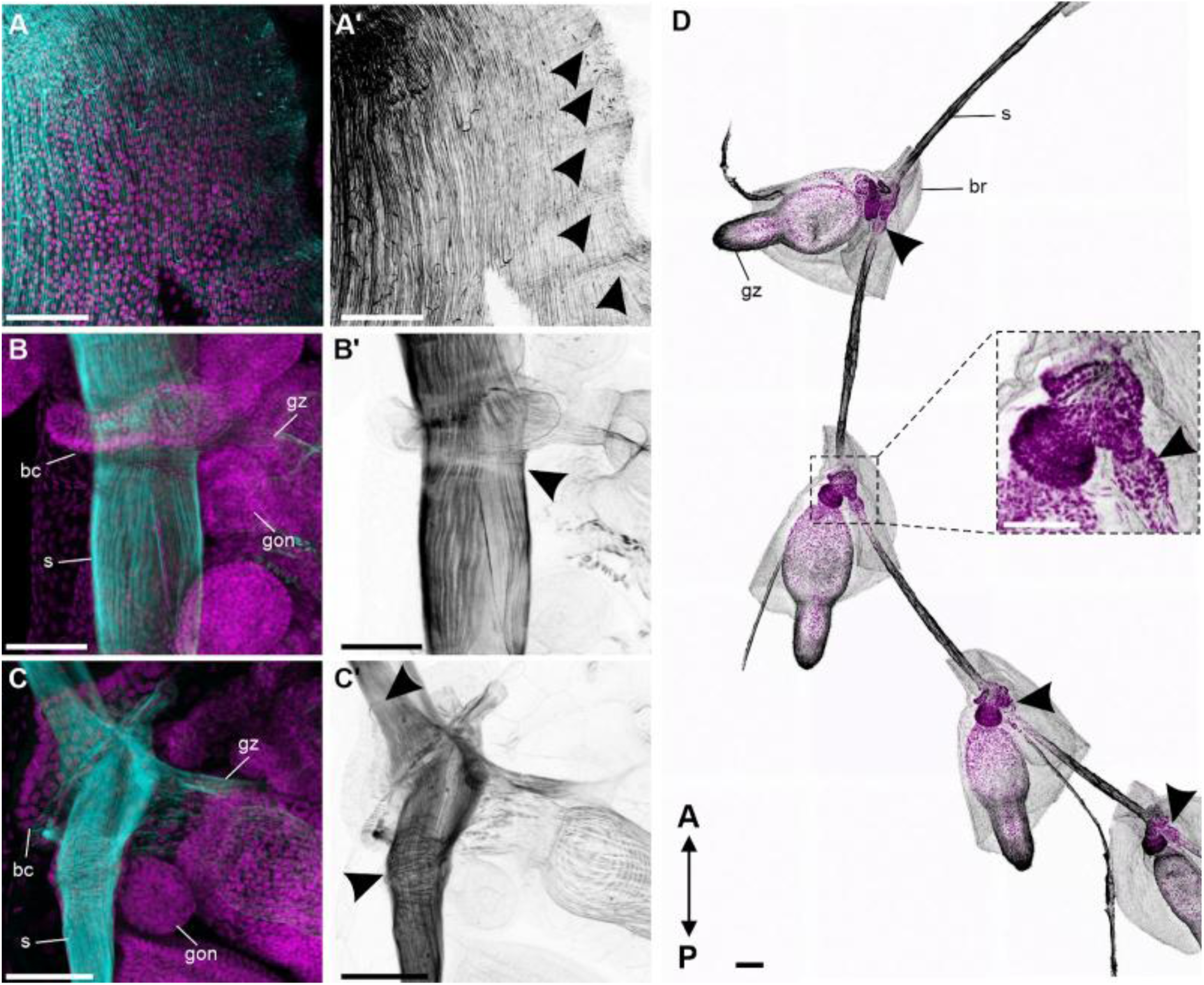
Detachment ring and stem-bracteal canal connection change during cormidium maturation. Actin (cyan in A, B, C, grey in A’, B’, C’) and nuclei (magenta) detection in the colony of *C. appendiculata.* **A**. A’. growth zone, **B. B’**. S2 cormidium, **C. C’**. S3 cormidium. Arrowheads point at the developing detachment rings (A, B, C). **D.** Tyrosinated tubulin (gray) and nuclei (magenta) detection in *C. appendiculata* stem with four S3 cormidia with a close-up of one of detachment rings (inset). Arrows point at detachment rings. A-P depicts anterior and posterior axis in all figures. Labels: bc – bracteal canal, br – bract, gon – gonophore, gz – gastrozooid, s - stem. Scale bars: 100 μm.

### Cormidium release induces eudoxid maturation

The process of release triggers eudoxid maturation, during which the bract undergoes a distinct morphological changes (Fig. 5). We again used time lapse videos to reconstruct the course of post-release maturation (Fig. 5A, Supplementary Movies 1-3). The newly released cormidium initially displayed an elongated stem fragment, which it fully resorbed within 4 h (Fig. 5A-G). As the stem resorbed, the bracteal canal elongated into the bract mesoglea, gradually narrowing down to form eudoxid’s phyllocyst – a swollen branch of bracteal canal, acting as nutrient reservoir (Fig. 5). This process occurred simultaneously with a shift in the anterior-posterior (A-P) axis of the eudoxid. In the *C. appendiculata* colony, the stem defines the A-P axis, positioning the bracteal canal perpendicularly to it (Fig. 2). During cormidium maturation, the apical projection of the bracteal canal develops at the angle to the colony’s A-P axis (Fig. 2A-E). Following release, the longer axis of the eudoxid’s phyllocyst becomes aligned with the A-P axis of the eudoxid (Fig. 5A-H).

**Fig. 5.**
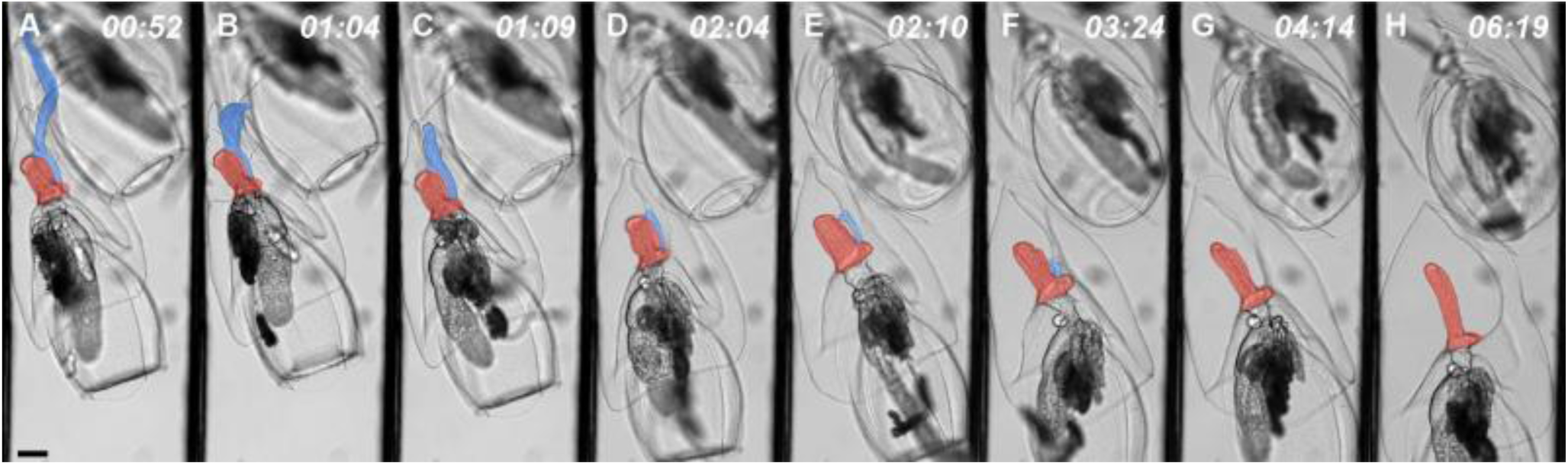
Eudoxid maturation comprises stem resorption, phyllocyst formation and bract remodeling. Individual frame grabs (**A-H**) with their timestamps (in hours after dissection) from time-lapse video depicting post-release modifications to cormidium morphology. Stem and bracteal canal/phyllocyst colored as in Fig. 2. Scale bar (shown only in A): 100 μm.

The most striking feature of eudoxid maturation is the remodeling of the bract, characterized by a significant increase in its volume (Fig. 5A-H). This process is likely accompanied by an uptake of water into the bract mesoglea rather than active growth, as our experiments were conducted on 1-day-starved specimens in microfiltered seawater. The resulting increase in volume, caused the bracteal furrow to close, which in turn led the bract’s shape to change from the thin, leaf-like structure wrapped around the stem into a pyramidal one positioned on top of the eudoxid.

### Eudoxid maturation is actomyosin dependent

Following cormidium release, the stem undergoes a series of contractions (Fig. 5A-C), likely facilitating its resorption (Fig. 6A, B). A strong actin signal at the connection site between the stem and the bracteal canal (Fig. 6A, B) indicates that it is an active site for the pulling of the stem remnant. In a fully mature eudoxid, the connection between the stem and bracteal canal was no longer visible, and the base of phyllocyst was composed of circularly arranged actin-rich structures of unknown function (Fig. 6C; Supplementary Fig. 1).

**Fig. 6.**
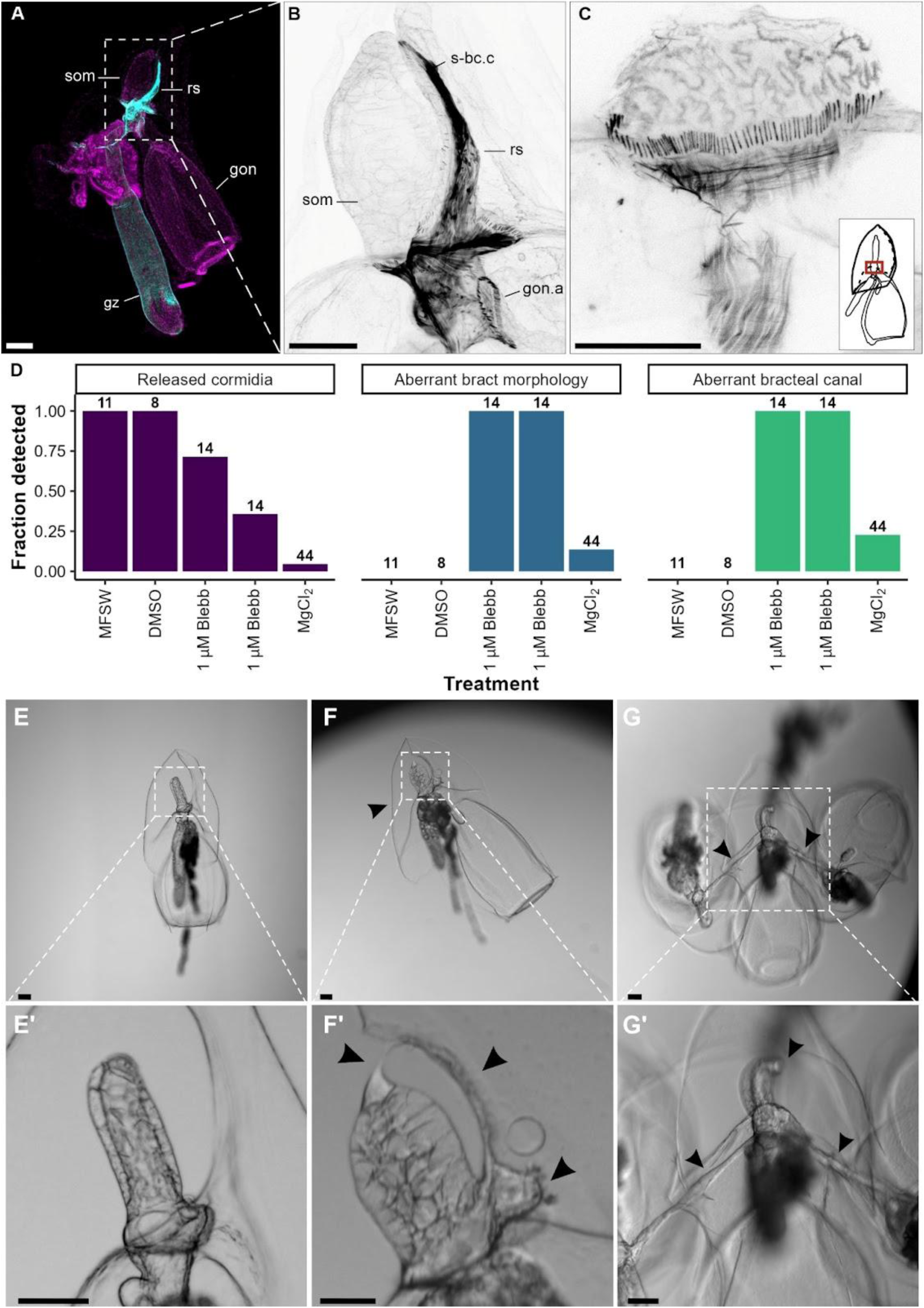
Cormidium release and eudoxid maturation depend on muscle activity. **A.** Actin (cyan) and nuclei (magenta) detection in newly released cormidium. **B.** Close up of A. depicting actin detection in stem-bract connection site. **C.** Actin detection at the base of mature eudoxid phyllocyst – red rectangle on the inset marks the location of the close up. **D**. Results of a pharmacological inhibition experiment scoring released cormidia, aberrant bract morphology and aberrant bracteal canal morphology across treatments with number of observations per treatment shown in bold (Supplementary Table 3; MFSW - microfiltered sea water, DMSO, 1μM Blebb - Blebbistatin in DMSO, 5μM Blebb - 5μM Blebbistatin in DMSO, isotonic MgCl_2_-isotonic 7.5% MgCl_2_ solution in MFSW). **E.** Normal phenotype of eudoxid from control MFSW treatment. **E’.** Close up of E showing phyllocyst morphology. **F.** Eudoxid with flattened bract (arrow) from 5μM Blebbistatin treatment. **F’.** Close up of F showing un-resorbed stem (right-hand arrows) and aberrant phyllocyst, connected with bracteal furrow (arrow on the left side). **G.** Three cormidia from MgCl_2_ treatment remaining interconnected with narrow stem (arrows), each with remodeled bract and some aberrations to phyllocyst formation. **G’**. Close up of G with arrows pointing at narrow stem (left and right arrow) and bent phyllocyst (top arrow). Scale bars:100 μm, except for C where scale bar is 50 μm.

In contrast, the absence of actin detection on the bract’s surface, before and after remodeling, along with a faint actin signal within the bracteal canal (Fig. 6A, B), imply that the bract remodeling is not driven by spatial differences in the tension of its surface epithelium or the bracteal canal. To assess if in turn myosin-dependent processes play a role in the bract remodeling, we revisited the previously mentioned pharmacological inhibition experiments and scored the resulting phenotypes (Fig. 6D-H, Supplementary Table 3). Treatment with magnesium chloride had no significant effect on bract remodeling, with only a small proportion of remodeled bracts displaying aberrant morphology (6/44; Fig. 6D). Similarly, the development of the bracteal canal into a phyllocyst appeared unaffected, with only 10 out of 44 cases showing a retained apical connection with bract’s surface. Interestingly, although cormidium detachment was not observed in magnesium chloride-treated 3-CLSF, the three cormidia still progressed through all stages of cormidium development and eudoxid maturation, but would remain interconnected by a narrow stem (Fig. 6D, H).

Treatment with the non-muscle myosin inhibitor, Blebbistatin, resulted in malformed bracts and bracteal canals (N = 28/28; Fig. 6D). Bracts in Blebbistatin-treated stem fragments continued to grow but failed at remodeling, which resulted in their pronounced flattening (Fig.6F). Bracteal canals expanded, similar to progression from S3 to S4 of cormidium development, but failed to elongate, rather becoming more oval. In addition, in released cormidia, the bracteal canal remained connected to the bract’s surface via the apical connection, causing it to bend towards the direction of the bracteal furrow (Fig. 6F, G). The stem resorption was also affected, as remnants of stem could have been discerned both along the bracteal furrow and near the bracteal canal base (Fig. 6G). Taken together, results of these experiments suggest that both bract remodeling and bracteal canal maturation into a phyllocyst – two key steps in eudoxid formation – are regulated by non-muscle myosin, inducing a massive reorganization of the region connecting the stem and the bracteal canal.

### A single acquisition of the eudoxid in calycophoran siphonophores

Our phylogenetic analyses provided a comprehensive picture of the evolutionary relationships within Calycophorae, thanks to the acquisition of new 16S and 18S sequences for previously unsampled species and genera (Fig. 7A, Supplementary Table 5). This robust phylogeny, constrained on a phylogenomic backbone^30^, allowed us to revise calycophoran families. We propose to dismiss the family Abyliidae L. Agassiz, 1862, as it is nested within Diphyidae Quoy & Gaimard, 1827. Since the family Clausophyidae Bigelow, 1913 appears polyphyletic, we propose to split it into two groups based on whether the genera produce eudoxids or not: (1) Chuniphyidae Moser, 1925, which contains the type genus *Chuniphyes*, as well as the other eudoxid-producing genera *Crystallophyes*, *Heteropyramis,* and *Kephyes*; and (2) the revised family Clausophyidae Bigelow, 1913, now including only one genus, *Clausophyes*, which does not produce eudoxids. The new clade Eudoxida, defined by the presence of a eudoxid in the life cycle, includes the families Chuniphyidae, Sphaeronectidae Huxley, 1859, and Diphyidae. The inclusion of *Tottonophyes enigmatica* in this clade remains ambiguous, awaiting confirmation that it produces eudoxids^31^.

**Fig. 7.**
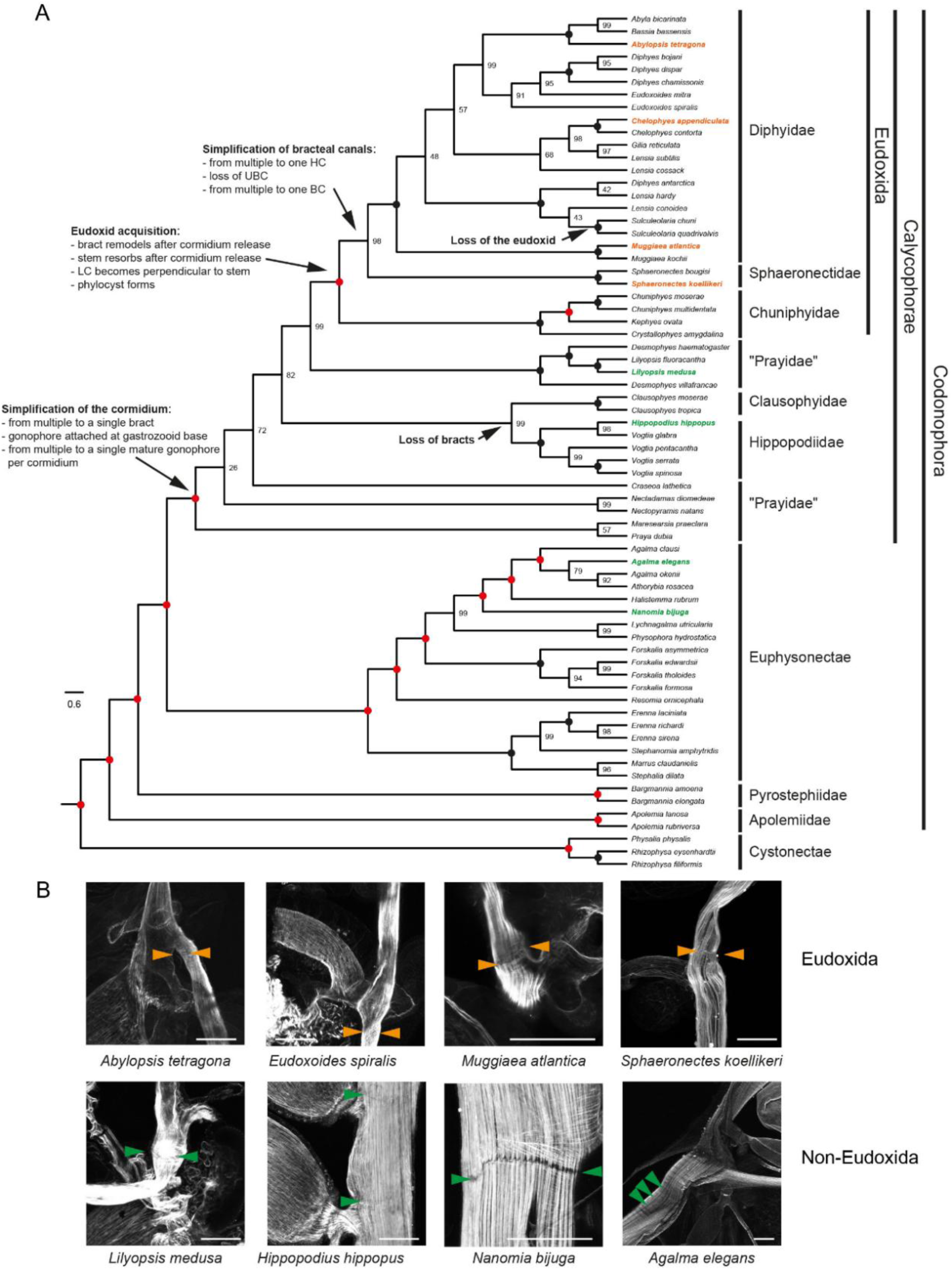
Siphonophore phylogeny based on 16S and 18S rDNA supporting a single origin of the eudoxid. **A.** ML phylogeny inferred from concatenated 18S and 16S sequences, constrained by the phylogenomic tree from Munro et al.^30^. Morphological evolution related to eudoxids is indicated on the phylogeny based on parsimony analysis. B. Actin (grey) detection in the stem for selected species shows a detachment ring in Eudoxida species (orange arrowhead), while a ‘break’ in the longitudinal fibers is observed at the corresponding location in non-Eudoxida species (green arrowhead). Scale bars: 100 μm.

By reconstructing 26 characters on the siphonophore phylogeny (Supplementary Data 1), we inferred that the eudoxid was acquired once in the common ancestor of Eudoxida, following a stepwise truncation of cormidium complexity and subsequent simplification of bract’s internal morphology (Fig. 7A). We also inferred a single loss of the eudoxid, in the ancestor *Sulculeolaria*, and loss of the bract in the common ancestor of Clausophyidae and Hippopodiidae. The ability of some prayid species to fragment their stem could not be linked to the origin of eudoxids, as this trait was found scattered among non-Eudoxida calycophorans.

The common ancestor of Codonophora possessed all the essential eudoxid building-blocks (bract, gastrozooid, gonophore), though some were present in more types or numbers than in Eudoxida (bracts, gastrozooids), others were arranged in more complex patterns (*e.g*., gonophores sharing a common pedicel) and additional ones, absent from extant Eudoxida, were present (palpons, Supplementary Data 1). Pronounced cormidium complexity decrease occurred in the common ancestor of calycophorans (Fig. 7A), marked by the presence of only one bract per cormidium. This was then followed, in Eudoxida, by an acquisition of traits likely involved in maintaining integrity (shift in bract attachment, stem resorption) and functional independence (*e.g*., phyllocyst) in released cormidia. A subsequent reduction in complexity led to gastrovascular canal system simplification within Diphyidae and Sphaeronectidae (Fig. 7A).

Cormidium release requires the presence of a detachment ring (Fig. 7B), which we inferred to be a synapomorphy of the clade Eudoxida. We could detect a detachment ring in the stem of five Eudoxida species (Fig. 7B, *C. appendiculata*: Fig. 3), while a ‘break’ in the longitudinal fibers was observed at the corresponding location in four non-Eudoxida codonophorans (Fig. 7B). Further investigations across Calycophorae are needed to infer the presence of the detachment ring in the common ancestor of Eudoxida.

### Spatiotemporal separation of eudoxids and colonies

Using extensive literature review we found evidence for distributional differences of eudoxids and polygastric colonies, across species and scales (temporal, horizontal and vertical; Fig. 8; Supplementary Data 2), with about 43% of references supporting that claim. Vertical distribution of eudoxids differed from that of polygastric colonies in a species-specific pattern, with either broader (*Abylopsis tetragona*^32^) or narrower (*Muggiaea bargmanna*^33^) depth-ranges of eudoxids, or their predominance at shallower (*Gilia reticulata*^34^) or great (*Diphyes antarctica*^34^) depths. Eudoxids also exhibited a distinct diel distribution pattern from polygastric colonies (Fig. 8A, E), suggesting differences in vertical migration, with one stage performing more pronounced migration than the other^40^.

**Fig. 8.**
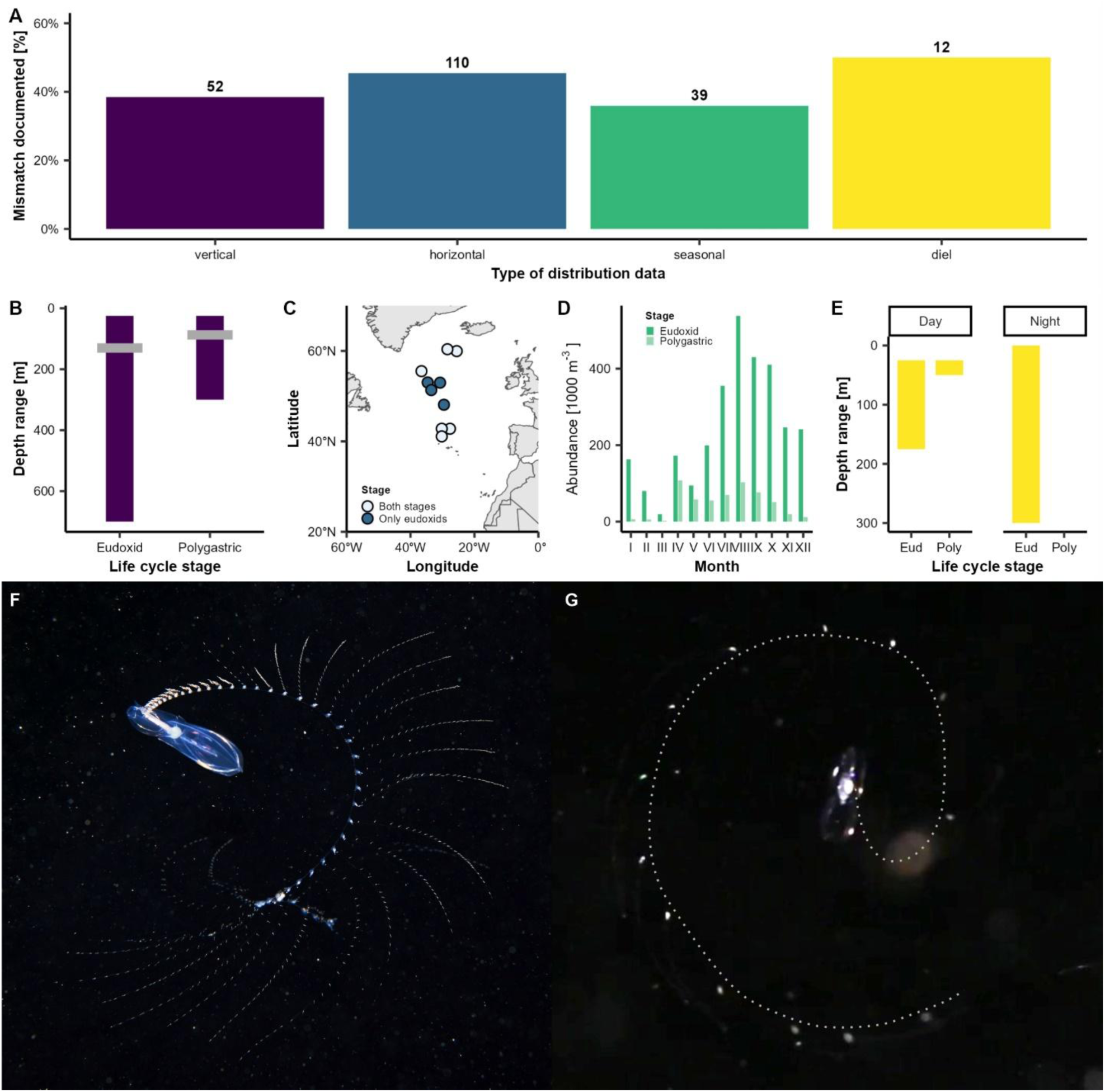
Ecological mismatch between eudoxid and polygastric life cycle stages. **A.** Results of literature review indicating percent of all references analyzed (numbers in bold; Supplementary Data 2) containing evidence for distribution mismatch between two life cycle stages separated into four types of distributional data: vertical, horizontal, seasonal and diel. **B.** Example of vertical distribution mismatch in *Abylopsis tetragona*, based on data from Grossmann et al.^32^, depicting depth ranges and mean distribution (grey line) of two life cycle stages. **C.** Example of horizontal distribution mismatch in *Chuniphyes multidentata,* based on data from Hosia et al.^36^, depicting sites where either both life cycle stages (light blue) or only eudoxids (dark blue) were present. **D.** Example of seasonal mismatch in eudoxid (dark green) and polygastric colonies (light green) of *Dimophyes arctica,* based on data from Hosia and Båmstedt^38^. **E.** Example of diel mismatch in distribution ranges of eudoxids (Eud) and polygastric colonies (Poly) in *Lensia cossack,* based on data from Grossmann et al.^32^. **F.** Photograph of *Chelophyes appendiculata* performing veronica movement captured by Alexander Semenov (White Sea Biological Station). **G.** Frame grab from video (Supplementary Movie 4) documenting *C. appendiculata* eudoxid performing veronica movement – dotted grey line indicates extent of eudoxid tentacle to facilitate interpretation.

The differences in vertical distribution of eudoxids and polygastric colonies could stem from distinct buoyancy of mature eudoxids, as compared to undetached cormidia. To test this hypothesis, we designed an experiment in which stem of a starved colony, spanning all S3 and S4 cormidia (five replicates, total N_cormidia_ = 89) was dissected off and transferred to a measuring cylinder filled with microfiltered sea water, where it immediately sank to the bottom (T_0_). We then recorded vertical position of individual cormidia, both at T_0_ and after 24h (Supplementary Table 5). We found that within that period all cormidia became released and matured to eudoxids, while simultaneously gaining positive buoyancy, likely through pronounced volume increase of the bract (Fig. 5).

We also found evidence of significant horizontal distribution mismatch between eudoxid and polygastric colonies, with eudoxid being present despite the absence of polygastric colonies (Fig. 8A, D). This was documented both in local studies (*Ceratocymba leuckartii*^35^) and those spanning greater oceanic regions (*Chuniphyes multidentata*^36^ Fig. 8C). These distributional differences often mirrored hydrological (*Heteropyramis maculata*^37^) and bathymetric (*Ceratocymba leuckarti*^35^) setting of the area, thus corroborating distinct environmental preferences of eudoxids and polygastric colonies. The lack of spatial overlap could also be indicatory of a temporal mismatch. Although the fact that eudoxids are produced by polygastric colonies, implies tight temporal coupling of their distributions, we found evidence to the contrary (Fig. 8A, D). For example, in a year-long study, presence of *Dimophyes arctica*^38^ eudoxids during periods of minimal or no presence of polygastric colonies, suggests extended lifespan of eudoxids and the ecological mismatch between the two.

### Eudoxids and colonies share complex foraging behavior

Despite differences in distribution, eudoxids share morphological traits with undetached cormidia, suggesting that their diets are likely similar to that of entire colonies. In most calycophoran polygastric colonies, feeding occurs through veronica movement^39^ in which nectophores pump to swim in circles, while stem and tentacles extend to form a feeding spiral that maximizes prey capture area (Fig. 8F). We observed similar behavior in eudoxids (Fig. 8G, Supplementary Movies 4-5), despite being achieved with different zooids. We documented for the first time *Chelophyes appendiculata* eudoxids mimicking foraging behavior of polygastric colonies, swimming in circles with gonophore pumping, simultaneously extending their tentacles.

## Discussion

Here, we describe the acquisition of an evolutionarily unique stage in the life cycles of colonial siphonophores, known as eudoxids. We provide evidence for the integration of zooids within a eudoxid, an evolutionary trajectory marked by a decreasing complexity of cormidia and zooids, and the acquisition of eudoxid-specific structures and processes (*e.g*., detachment ring, phyllocyst, post-release maturation). Additionally, we documented distributional niche partitioning between the calycophoran life cycle stages, and the mimicking of colony’s complex foraging behavior by eudoxids, resulting from the cooperative actions of zooids. Thus, we consider eudoxids to exhibit all hallmarks of individuality: cooperative, functional, and evolutionary^3,20^.

### Evolution of siphonophore colony architecture

In agreement with Dunn and Wagner^12^, we show that calycophoran cormidia are more modular than in other siphonophores, in that individual zooids are not dispersed along the stem, but rather remain clustered. Our data, however, indicate that *Chelophyes appendiculata* budding bypasses the probud subdivision, in that the original bud develops directly into a gastrozooid, while subsequent zooids are budded directly from gastrozooids’ base (Fig. 2A). Alteration of probud subdivision might be a shared feature of Calycophorae, as it was indicated in two other species^40,41^. Future exploration of cormidium budding across calycophorans, especially among “prayid” siphonophores, will be necessary to better understand the evolution of siphonophore colony architecture.

The apparent increase in cormidia modularity in Calycophorae coincided with the loss of their complexity (Fig. 2, 7). Aside from the reduction in zooid types and numbers, we also documented the alteration to the bract morphology, following eudoxid acquisition (Fig. 7). This change involved twisting the bract attachment to the stem through the longitudinal canal lamella, resulting in the canal becoming perpendicular to the stem (Fig. 7, Supplementary Data 1). Given the likely muscular nature of these lamellae^18,42^ (Fig. 2C, 3E, 6C, Supplementary Fig. 1), such repositioning might have secured a stronger attachment of the bract, necessary for mature eudoxid functioning and complex behavior (Fig. 8G). The subsequent evolutionary modifications of the bract included a reduction in the number of bracteal canal branches (Fig. 7, Supplementary Data 1) leading to the retention of only the one forming the phyllocyst after cormidium detachment.

### Mechanisms of siphonophore colony fragmentation

The anterior-posterior patterning of the linear colonies in siphonophores, and their directional growth with strongly localized budding and stem cell populations^43^, hampers module replacement and removal, as indicated by the limited regenerative potential of siphonophores^4,39,44^. One plausible outcome of the limited regenerative potential of siphonophores is the phylogenetically broad distribution of records of colony fragmentation^45^ (Fig. 7, Supplementary Data 1). Some siphonophore colonies attain enormous lengths, notably apolemiid “physonects” (over 30 m^18,45^) and “prayid” Calycophorae (45.7 m in *Praya dubia*^46^), and stem fragmentation was documented in these two groups (apolemiids^47^, “prayids”^26^; Supplementary Data 1). These freely drifting stem fragments vary in length, module composition and reproductive capacity^18,47^, indicating the unprogrammed nature of their separation, likely driven by the injury to the colony. In contrast, we show here that colony fragmentation leading to eudoxid formation (*i.e*., cormidium release and eudoxid maturation) is a precisely timed process, relying on module (here cormidium) development (Fig. 4, Supplementary Table 1-2) and loss of connectivity with the colony (Supplementary Table 2).

The temporal and spatial separation of eudoxid and cormidium maturation, suggests presence of an inhibitory signal suppressing the bract remodeling, phyllocyst formation and stem resorption before cormidium release (Fig. 5). The nature of this signal remains to be determined, but our data on the detachment ring role in the cormidium release (Fig. 3A, B, D) and on the cormidium release from stem fragments (Supplementary Table 2), suggests that it is the loss of connectivity with the colony that induces cormidium release and eudoxid maturation. It is likely, given the complex coordination system present in *C. appendiculata* and other siphonophores^48^, that sensing of cormidium integration with the colony (presence of connectivity) may involve the nervous system. Supporting this claim, we documented the existence of transverse neuronal bands in stems of *C. appendiculata* (Fig. 2A), that mirrored the position of detachment rings and remained connected with the two giant axons running along the entirety of the stem (Fig. 2A, Supplementary Fig. 2A), supporting this claim. Alternatively, sensing of connectivity may be related to the flow in the stem^48^, food availability or a mix of both. Our observations on spatial distribution of the detachment rings (Fig. 4D) and their constriction prior to the detachment (Fig. 3) suggests, that they play the role in restricting flow across the stem, potentially serving as a cue to initiate cormidium release. Interestingly, while we observed detachment rings only in siphonophore species releasing eudoxids (Fig. 7), in non-Eudoxida we observed presence of transverse actin-rich bands in corresponding sites (Fig. 7), consistent with earlier studies that hypothesized their role in controlling local stem contractions^42^. This may indicate that eudoxid-releasing calycophorans have co-opted neuro-muscular toolkit present in other siphonophores, using detachment rings to mediate intra-colony connectivity.

### Role of muscles in eudoxid production

Through multiple lines of evidence (Fig. 4, 6), we found that cormidium release depends on muscle contraction of the detachment ring, likely a Eudoxida-unique structure (Fig. 7). Despite functional diversity of cnidarian muscles^49^, information on their involvement in reproductive processes and life cycle transitions is scarce. Neuro-muscular control of zooid detachment was suggested during shedding of bracts and nectophores^42^, as well as reproductive release of gonophores^18^. Those three zooids share a similar structure of the actin-rich stem attachment at their base, called the lamella or “autotomy joints”^18,42^ (Fig. 2C, 3E, 6C, Supplementary Fig. 1), which could be contractile and participate in zooid detachment. Additionally, bud detachment in *Hydra* was shown to rely on actin dynamics and regionally specific myosin phosphorylation for parent-bud boundary formation and detachment by constriction^50^, and this resembles, to some extent, constriction-dependent eudoxid release (Fig. 4). We could show that eudoxid maturation is controlled by actomyosin (Fig. 6), reminiscent of the role of non-muscle myosin in the remodeling of regenerating *Clytia hemisphaerica* jellyfish^51^ and in affecting growth of *Podocoryna carnea* stolons^52^.

### Ecological success of eudoxids

Fishing postures in siphonophores are closely related to their trophic niche^53^. The so-called veronica movement, observed in both eudoxids and polygastric colonies (Fig. 8F, G), appears to be restricted to members of Eudoxida. In contrast, other siphonophores, particularly those with longer and less contractile stems, adopt a “long-line” fishing posture^18,53^. The mechanisms underlying the acquisition and coordination of eudoxid behavioral complexity remain to be determined. One possibility is that it involves nervous system rearrangement. Indeed, we found that the two giant axons of the stem disappear during stem resorption (Supplementary Fig. 2A, B).

The common assumption regarding the distributions of eudoxids and parental colonies, is that both co-occur within the same environment^27,54^. Through a comprehensive literature review we found evidence challenging this widely held view (Fig. 8, Supplementary Data 2). Of 386 references related to calycophoran distribution, we identified 122 studies that counted abundances of both life cycle stages separately, and among these 43% documented distributional mismatches (horizontal, vertical, seasonal, diel) between eudoxids and polygastric colonies (Fig. 8, Supplementary Data 2). These mismatches may help reduce intraspecific competition – a likely concern for calycophorans, given that veronica movement in both life cycle stages of Eudoxida suggests similar dietary preferences (Fig. 8). This is consistent with data from other marine taxa with complex life cycles, which show that differences in habitat use, trophic modes and mobility across life stages are key drivers of ecological success^11,55^.

## Materials and Methods

### Collection and maintenance of *Chelophyes appendiculata*

Live specimens of *Chelophyes appendiculata* used in all experiments were collected in spring (2018-2022) from the Villefranche Bay (northern Mediterranean Sea), either in the vicinity of the Point B (43.6830 N, 7.3170 E) monitoring station or near the entrance of the bay. Sampling occurred in the early morning hours onboard RV Pelagia (Institute de la Mer de Villefranche-sur-Mer, France). To fish siphonophores, a plankton net (650 µm mesh) was dropped below the surface (1-2 m) and was slowly towed behind the vessel to avoid damage to specimens. Then, the cod-end was gently emptied into a 15L bucket (CAMBRO ®) prefilled with natural seawater. In case of large concentrations of plankton (*e.g.,* during salp bloom) colonies of *C. appendiculata* were immediately transferred to a separate bucket with ladles.

After collection, specimens were moved to a temperature-controlled room (18°C) with a 12h:12h light cycle. Colonies designated for experiments were maintained unfed, in 15L buckets filled with filtered sea water, for no longer than 3 days. All other specimens (*e.g.*, for immunostaining) were kept in 17L Kreisel tanks (Exotic Aquaculture) for 1-2 weeks, were fed daily with a mix of zooplankton, collected with 180 μm mesh plankton net from the Villefranche Bay and were starved only 1 day prior to use.

### Initial processing of specimens

Specimens designated for subsequent analyzes were individually placed in a Petri dish coated with Sylgard 184 (Dow Corning Corporation) filled with microfiltered sea water (MFSW) and relaxed by dropwise addition of isotonic 7.5% solution of magnesium chloride hexahydrate in Mili-Q water at 1/3 MgCl_2_ to 2/3 MFSW ratio. As tentacles tended to entangle and thus obscure observations, they were trimmed with an angled-micro knife (22.5°; Fine Science Tools GmbH). When stem fragments were needed, they were dissected from colonies also using angled micro-knife.

### *In vivo* experiments

Ethics approval is not required for experimentation on cnidarians. The following *in vivo* experiments were run on 1-day starved *C. appendiculata*: Competence to release (Supplementary Table 1), Stem fragmentation (Supplementary Table 2; Supplementary Movies 1-3), Buoyancy (Supplementary Table 6) and Eudoxid behavior (Supplementary Movies 4-5). They were all run in a temperature-controlled room (18°C).

#### Competence to release

*Chelophyes appendiculata* colonies (nectophore and entire stem) used were initially relaxed (as described above) in order to stage and photograph (NEX-5R Sony camera mounted on an Olympus SZ61 stereomicroscope) the most-posterior cormidium. To avoid specimen damage due to tentacle entanglement all tentacles were trimmed. After staging, colonies were transferred to one of two treatments: unperturbed or immobilized. In unperturbed treatment, specimens were transferred to 1L glass beakers filled with MFSW. In case of immobilized treatment, specimens were individually placed in Sylgard 184 coated Petri dish, prefilled with solution of 7.5% magnesium chloride hexahydrate in Mili-Q mixed with MFSW in 1:2. The anterior nectophore of a colony was then pinned down to Sylgard using ethanol-washed Opuntia spines. Usually, three spines were used per nectophore, with one driven through nectophore tissues near the ostium, and two placed on both sides of the anterior tip of the nectophore, which successfully limited nectophore movement. Then, the magnesium chloride solution was gently replaced with MFSW with antibiotics (1/2000 dilution of a penicillin [10000 units stock] and streptomycin [10 mg per ml] solution; Sigma Aldrich, #P4333), which was replaced daily. The experiments were run for either 24h or 48h, with observations every 2-3h, except for nights. Upon every release event, the stage of released cormidium was noted down (Supplementary Table 1).

#### Stem fragmentation

Stems of *C. appendiculata* colonies were relaxed and their tentacles trimmed as described above. They were then cut into 2-, 3- or 4-cormidia long stem fragments (once a single cormidium was dissected off, Supplementary Movie 3), and the stage of development of individual cormidia was noted down. The fragments were then individually transferred either into 6-well plates or glass micro-chambers. In case of 6-well plates, they were prefilled with 6 ml of MFSW with antibiotics (as above; replenished daily), and fragments were observed Olympus SZ61 stereo-microscope with NEX-5R Sony camera at varying frequency (Supplementary Table 2), with the picture taken at every time point and observations of cormidium release noted down. The glass micro-chambers were made by breaking glass microscope slides and positioning resulting pieces in a Petri dish (3 cm diameter) to form a channel, roughly 1 mm in width. The chambers were then placed directly under Zeiss Axio Observer, prefilled with MFSW with antibiotics to the upper level of glass slide upper surface, and the stem fragment was put directly to the channel. The movement of the stem fragment within the channel was then constrained by placing Opuntia spines on either side of the channel. Zeiss’s ZEN Microscopy Software was then used to set automatic picture capture every 1 or 2 minutes, for about 24h, across the full z-extent of the channel. To minimize file size, these pictures were then exported as movies (Supplementary Movies 1-3) at high frames-per-second rating and at a single z-plane, chosen based on stem fragment position throughout the experiment.

#### Buoyancy

Five experiments were run, each with a single *C. appendiculata* colony. Each colony was first relaxed (as described in the Initial processing of specimens section), all tentacles were trimmed, and a single stem fragment was dissected off that spanned all S3 cormidia. The stem fragment was then placed in a 50 ml measuring cylinder, prefilled with MFSW with antibiotics (as described above), and the position of cormidia was recorded as either laying on the bottom (0) or floating (1). Then the subsequent recording occurred after 24 h (Supplementary Table 5).

#### Eudoxid behavior

Twenty eudoxids were taken from Kreisel tank where polygastric colonies were kept and were transferred to a 1L rectangular tank filled with MFSW. Eudoxid behavior was recorded using Canon EOS 6D camera with Canon 100 mm f/2.8 L EF Macro IS USM lens (Supplementary Movies 4-5).

### Immunostaining and actin detection

Specimens designated for immunohistochemistry were processed as in Mańko et al.^14^ using anti-tyrosinated tubulin antibody (YL1/2, 1:50, Abcam, #6160) and/or anti-FMRFamide antibody primary antibodies coupled with secondary antibodies, Alexa Fluor 488-conjugated phalloidin (1:50, solubilized in methanol; Thermo Fisher Scientific, #10125092), and Hoechst (Sigma Aldrich, #94403, 1/2000). Samples were gradually (10%-20%-40%) transferred to Citifluor AF-1 antifade mountant for imaging. Preparations were then visualized under Leica’s SP5 and SP8 confocal microscopes. Given the large size of imaged animals, multiple tiles spanning all-z planes were often taken. These were automatically aligned using Leica Application Suite X Life Science Microscope software, while z-stacks were projected using standard deviation projection in ImageJ.

### Pharmacological inhibition experiments

2- and 3-cormidia long stem fragments (Supplementary Table 3) were individually transferred to 6-well plates prefilled with 10 ml of solutions corresponding to treatments analyzed. These included: 1 or 5 µM Blebbistatin (Sigma-Aldrich, #B0560; solution in MFSW, diluted from a 34 mM stock solution in DMSO); MgCl_2_ (solution of 7.5% magnesium chloride hexahydrate in Mili-Q mixed with MFSW in 1:2 proportion); DMSO (1.47 µL DMSO in MFSW, corresponding to DMSO concentration in 5 µM blebbistatin treatment); MFSW. Experiments were run in temperature-controlled conditions at 18°C, and antibiotics were added to each treatment as described above. The development stage of each cormidium was staged in each stem fragment analyzed and individually photographed either under Zeiss Axio Observer or using Sony camera (NEX-5R) mounted on an Olympus SZ61 stereomicroscope. They were imaged again after 22h and their phenotype was scored in the following categories: released cormidia (number of cormidia released from stem fragment); remodeled bracts (number of bracts that whose morphology matches that of released eudoxid bract); aberrant bract morphology (number of bracts whose morphology differs from that of eudoxid bract); aberrant bracteal canal (number of bracts showing alteration of bracteal canal morphology); stems resorbed (number of cormidia not showing signs of presence of loose stem).

### Electron microscopy

Two samples (3-cormidia long stem fragment and two eudoxids of *C. appendiculata*) were preprocessed (relaxing and dissections) as described in Initial processing of specimens section. Specimens were then transferred separately to 1.5 ml Eppendorf tube and were fixed overnight at 4°C in a solution of 0.05M sodium cacodylate buffer (pH 7.2-7.4), 0.025% MgCl_2_, 2.5% glutaraldehyde, 0.8% paraformaldehyde and 4.45M NaCl in deionized water. Subsequent steps leading to resin embedding were conducted at the Plateforme Commune de Microscopie Electronique, Université Nice Côte d’Azur. Transmission electron microscopy images of ultrathin sections were acquired on Tecnai G2 Spirit BioTWIN (FEI) at 120 kV at the Electron Microscopy Section at the Faculty of Biology at the University of Gdańsk.

### DNA Extraction & sequencing

DNA was extracted from nectophores or a portion of the siphosome stored in ethanol at -20°C. DNA was extracted using the DNeasy Blood & Tissue Kit (Qiagen), following kit instructions. As much ethanol was removed from samples as possible before adding buffer ATL. Samples were incubated at 56°C for 3 hours. PCR of 16S and 18S was done with the following primers (16S: SHA 5’ ACGGAATGAACTCAAATCATGT 3’, SHB 5’ TCGACTGTTTACCAAAAACATA 3’, 74; 18S MitchA 5’ AACCTGGTTGATCCTGCCAGT 3’, MitchB 5’ TGATCCTTCTGCAGGTTCACCTAC 3’, 75), using the Phusion high fidelity polymerase (New England Biolabs) and annealing temperatures of 50°C and 55°C respectively (for cycling profiles see 16S:33, 18S:76). PCR products were cleaned up using the QIAquick PCR Purification Kit (Qiagen) following kit instructions and were sent to Eurofins Genomics GmbH (Germany) for Sanger sequencing. In addition to using the same forward and reverse PCR primers for sequencing, the following internal primers were used for 18S sequencing: 18SR1028 5’CTGCGAAAGCATTTGCCAAG 3’ and 18SF970: 5’CTAGGACGGTATCTGATCGTCTTCG 3’,77). All new sequences generated here were deposited at GenBank (accession numbers: PV190272-PV190290; Supplementary Table 4).

### Phylogenetic analyses

Available 16S and 18S rDNA sequences of siphonophores were retrieved from NCBI, except for *Physophora gilmeri* and *Thermopalia taraxaca*, whose 18S sequences were partial, *and Rosacea flaccida* and *Gymnopraia lapislazula*, whose phylogenetic positions were poorly supported. Additionally, because the positions of *Cordagalma ordinatum* and *Frillagalma vityazi* in rDNA-based phylogenies^56,57^ were incongruent with those in phylogenomic-based phylogenies^30^, these species were also excluded from the analyses. Four species included in the molecular phylogeny (*Dendrogramma enigmatica, Tottonophyes enigmatica, Sphaeronectes haddocki, and Sphaeronectes christiansonae*) were excluded from the analyses due to the lack of information about their eudoxids.

We added new sequences for 18 calycophoran species for both ribosomal markers, as well as the 18S of *Lilyopsis medusa*. The outgroup included five hydrozoan species: *Ectopleura dumortieri, Hydra circumcincta, Staurocladia wellingtoni, Porpita porpita*, and *Velella velella*. We aligned the sequences using MAFFT v7.271 L-INS-I algorithm^58^ and conducted constrained and unconstrained Maximum Likelihood (ML) analyses on the concatenated 16S-18S alignment using IQTree^59^ with 1000 bootstrap replicate. ModelFinder implemented in IQTree v1.5.5. was used to assess the relative model fit, selecting GTR+R4 for having the lowest Bayesian Information Criterion score. The constrained tree topology was derived from Munro et al.^30^ (Supplementary Fig. 5, Supplementary Data 4).

A matrix including 26 morphological characters coded for all described siphonophore species was assembled from an extensive literature review (Supplementary Data 1, 4). For each species included in the phylogeny, we first analyzed the original species description, and in case of absence of information on a given trait we subsequently checked information in morphological reviews (Supplementary Table 1). We then encoded additional information stemming from present work. The evolution of these characters was inferred by parsimony onto the ML phylogeny using Mesquite^60^.

### Siphonophore ecology review

Information on distribution mismatch between polygastric colonies and eudoxids were retrieved from a review of siphonophore reference database curated by late Philip R. Pugh (National Oceanographic Centre), including 1655 positions, spanning years 1725-2023. First, the type of each reference was verified, and only scientific papers and books were retained, removing technical reports, regional species checklists, theses, and grey literature. Then we inferred the field of study for each of those by analysis of the title and abstract, retaining only references containing information on species distribution (vertical, horizontal, seasonal, or diel; N=520). We then verified if these contain information on distribution of Calycophorae, excluding references dealing either collectively with siphonophores, or not mentioning calycophores (N=386). From these we then kept only those that include information on eudoxids (N=122, Supplementary Data 2), and these were then read thoroughly and checked for indication of distribution mismatch. Distribution was classified as pointing at mismatch when: for horizontal data, eudoxids present at sites when no polygastric stages were present, for vertical data, eudoxids showing different depth range or mean depth distribution than polygastric, for seasonal data, eudoxids not following seasonal abundance cycle of polygastric stages (e.g., presence in seasons when no or almost no polygastric stages were recorded), for diurnal data, eudoxids showing different vertical distribution between day and night as compared to polygastric stages.

Example plots were generated using raw data provided in the papers (*Chuniphyes multidentata* horizontal distribution^36^, *Abylopsis tetragona* vertical and *Lensia cossack* diel distributions^32^) or data were retrieved through digitization of published plots using plotdigitzer.com (*Dimophyes arctica* seasonal distribution^38^).

## Acknowledgments

We would like to thank numerous people who have made this work possible, by assisting us in sampling in the Villefranche Bay (Alexandre Alié, Institut de la Mer de Villefranche (IMEV), Alexandre Jan, University of Bergen), aiding in *C. appendiculata* maintenance (Alexandre Jan, University of Bergen), helping with confocal (Sébastien Schaub, IMEV; Michał Rychłowski, University of Gdańsk) and electron microscopy (Bastien Salmon, BIOM; Sophie Pagnotta, Université Nice Côte d’Azur; Dorota Łuszczek and Magdalena Narajczyk, University of Gdańsk), and to Alexander Semenov (White Sea Biological Station) for providing us with his photograph of *C. appendiculata,* Anna Panasiuk for providing samples of *Diphyes antartctica* and Samuel Church (Yale University) for commenting on earlier version of manuscript. MKM also thanks Steven Haddock (Monterey Bay Aquarium Research Institute) for the invitation to join research cruise to Central Pacific, where some of the animals were sampled, and the Laboratoire de Biologie du Développement at the (IMEV), Clytia team in particular, for their hospitality during his numerous visits. We are grateful to the Imaging Platform (PIM), animal facility (CRB) and sailors (Jean-Yves Carval, in particular) of the IMEV, which is supported by EMBRC-France, whose French state funds are managed by the ANR within the Investments of the Future program under reference ANR-10-INBS-0, for continuous support. We also acknowledge the Electron Microscopy facility CCMA (Centre Commun de Microscopie Appliquée) from the Université Côte d’Azur, part of the Microscopie Imagerie Côte d’Azur GIS IBiSA labeled platform, supported by Université Côte d’Azur, the Région Sud, and the Département 06.

This work received financial support from the following grants: National Science Centre in Poland, Preludium 15, 2018/29/N/NZ8/01305 (MKM, LL), University of Gdańsk, UGrant-START 533-O000-GS10-23 (MKM), European Marine Biology Resource Centre, ACCESS, OOV–EMBRC FR–AAP2018–2180 (MKM, LL).

## Author contributions

Conceptualization: M.K.M., L.L.

Methodology: M.K.M., L.L.

Investigation: M.K.M., L.L., C.M.

Formal analysis: M.K.M., L.L.,

Funding acquisition: M.K.M., L.L.,

Visualization: M.K.M.

Writing - Original Draft: M.K.M

Writing - Review & Editing: L.L., C.M.

Supervision: L.L.

## Data and materials availability

All data needed to evaluate the conclusions in the paper are present in the paper and/or the Supplementary Materials.

## References

1. Buss, L. W. The Evolution of Individuality. (Princeton University Press, Princeton, 1987).

2. Smith, J. M. & Szathmáry, E. The Major Transitions in Evolution. (Oxford University Press, 1995).

3. Carmel, Y. & Shavit, A. Operationalizing evolutionary transitions in individuality. Proceedings of the Royal Society B: Biological Sciences 287, (2020).

4. Mackie, G. O. From aggregates to integrates: physiological aspects of modularity in colonial animals. *Philosophical Transactions - Royal Society of London*, Series B 313, 175–196 (1986).

5. Cartwright, P., Travert, M. K. & Sanders, S. M. The evolution and development of coloniality in hydrozoans. J Exp Zool B Mol Dev Evol 336, 293–299 (2021).

6. Hiebert, L. S., Simpson, C. & Tiozzo, S. Coloniality, clonality, and modularity in animals: The elephant in the room. J Exp Zool B Mol Dev Evol 336, 198–211 (2021).

7. Brown, F. D. Evolution of animal coloniality and modularity: Emerging themes. J Exp Zool B Mol Dev Evol 336, 187–190 (2021).

8. Phillippi, A. L. & Yund, P. O. Self-fertilization and inbreeding depression in three ascidian species that differ in genetic dispersal potential. Mar Biol 164, (2017).

9. Burgess, S. C., Baskett, M. L., Grosberg, R. K., Morgan, S. G. & Strathmann, R. R. When is dispersal for dispersal? Unifying marine and terrestrial perspectives. Biol Rev Camb Philos Soc 91, 867–882 (2016).

10. Gemmell, B. J. et al. Cool your jets: biological jet propulsion in marine invertebrates. Journal of Experimental Biology 224, jeb222083 (2021).

11. Boosten, M. et al. Loss of the benthic life stage in Medusozoa and colonization of the open ocean. Preprint at 10.1101/2023.02.15.528668 (2023).

12. Dunn, C. W. & Wagner, G. P. The evolution of colony-level development in the Siphonophora (Cnidaria:Hydrozoa). Dev Genes Evol 216, 743–754 (2006).

13. Totton, A. K. & Bargmann, H. E. A Synopsis of Siphonophora. (Trustees of the British Museum (Natural History), Lodon, 1965).

14. Mańko, M. K., Munro, C. & Leclère, L. Establishing Bilateral Symmetry in Hydrozoan Planula Larvae, a Review of Siphonophore Early Development. Integr Comp Biol 63, 975– 989 (2023).

15. Munro, C., Vue, Z., Behringer, R. R. & Dunn, C. W. Morphology and development of the Portuguese man of war, *Physalia physalis*. Sci Rep 9, 1–12 (2019).

16. Travert, M. et al. Coevolution of the Tlx homeobox gene with medusa development (Cnidaria: Medusozoa). Commun Biol 6, 709 (2023).

17. Oderberg, D. S. Siphonophores. A metaphysical case study. in Biological Identity (eds. Meincke, A. S. & Dupré, J.) (Routledge, London, 2020).

18. Mackie, G. O., Pugh, P. R. & Purcell, J. E. Siphonophore Biology. Adv Mar Biol 24, 97– 262 (1987).

19. Mackie, G. O. Siphonophores, Bud Colonies, and Superorganism. in The Lower Metazoa (ed. Dougherty, E.) 329–337 (University of California Press, Berkeley, 1963).

20. Díaz-Muñoz, S. L., Boddy, A. M., Dantas, G., Waters, C. M. & Bronstein, J. L. Contextual organismality: Beyond pattern to process in the emergence of organisms. Evolution (N Y) 70, 2669–2677 (2016).

21. Müller, P. E. Iagttagelser over Nogle Siphonophores. (Jacob Lunds Boghandel, Kopenhagen, 1871).

22. Haeckel, E. Report on the Siphonophorae collected by HMS Challenger during the years 1873-1876. Report of the Scientific Results of the voyage of H.M.S. Challenger. Zoology 28, 1–380 (1888).

23. Schuchert, P. DNA barcoding of some Pandeidae species (Cnidaria, Hydrozoa, Anthoathecata). Revue suisse de Zoologie 125, 101–127 (2018).

24. Gegenbaur, C. Beitrage zur naheren Kenntniss der Schwimmpolypen. (Siphonophoren.). Wissenschaftliche Zoologie 5, (1853).

25. Leuckart, R. Zoologische Untersuchungen. 1. Die Siphonophoren. (J. Ricker’sche Buchhandlung, Giessen, 1853).

26. Totton, A. K. Siphonophora of the Indian Ocean together with systematic and biological notes on related specimens from other oceans. Discovery Reports 27, 1–162 (1954).

27. Grossmann, M. M., Lindsay, D. J. & Collins, A. G. The end of an enigmatic taxon: Eudoxia macra is the eudoxid stage of Lensia cossack (Siphonophora, Cnidaria). Syst Biodivers 11, 381–387 (2013).

28. Licandro, P., Souissi, S., Ibanez, F. & Carré, C. Long-term variability and environmental preferences of calycophoran siphonophores in the Bay of Villefranche (north-western Mediterranean). Prog Oceanogr 97–100, 152–163 (2012).

29. Haddock, S. H. D., Dunn, C. W. & Pugh, P. R. A re-examination of siphonophore terminology and morphology, applied to the description of two new prayine species with remarkable bio-optical properties. Journal of the Marine Biological Association of the United Kingdom 85, 695–707 (2005).

30. Munro, C. et al. Improved phylogenetic resolution within Siphonophora (Cnidaria) with implications for trait evolution. Mol Phylogenet Evol 127, 823–833 (2018).

31. Pugh, P. R., Dunn, C. W. & Haddock, S. H. D. Description of *Tottonophyes enigmatica gen. nov., sp. nov.* (Hydrozoa, Siphonophora, Calycophorae), with a reappraisal of the function and homology of nectophoral canals. Zootaxa 4415, 452–472 (2018).

32. Grossmann, M. M., Nishikawa, J. & Lindsay, D. J. Diversity and community structure of pelagic cnidarians in the Celebes and Sulu Seas, southeast Asian tropical marginal seas. Deep Sea Res 1 Oceanogr Res Pap 100, 54–63 (2015).

33. Pagès, F. & Kurbjeweit, F. Vertical distribution and abundance of mesoplanktonic medusae and siphonophores from the Weddell Sea, Antarctica. Polar Biol 14, 243–251 (1994).

34. Pugh, P. R., Pagès, F. & Boorman, B. Vertical distribution and abundance of pelagic cnidarians in the Eastern Weddell Sea, Antarctica. Journal of the Marine Biological Association of the United Kingdom 77, 341–360 (1997).

35. Rengarajan, K. Quantitative and seasonal abundance of siphonophores along the southwest coast of India and the Laccadive Sea. J. mar. bioI. Ass. India 25, 17–40 (1983).

36. Hosia, A., Stemmann, L. & Youngbluth, M. Distribution of net-collected planktonic cnidarians along the northern Mid-Atlantic Ridge and their associations with the main water masses. Deep Sea Res 2 Top Stud Oceanogr 55, 106–118 (2008).

37. Panasiuk, A., Grzonka, L., Prątnicka, P., Wawrzynek-Borejko, J. & Szymelfenig, M. Zonal variability of pelagic Siphonophora (Cnidaria) in the Atlantic sector of the Southern Ocean. J Sea Res 165, 101951 (2020).

38. Hosia, A. & Båmstedt, U. Seasonal abundance and vertical distribution of siphonophores in western Norwegian fjords. J Plankton Res 30, 951–962 (2008).

39. Mackie, G. O. & Boag, D. A. Fishing, feeding and digestion in siphonophores. Pubbl. Staz. Zool. Napoli 33, 178–196 (1963).

40. Schneider, K. C. Mittheilungen über Siphonophoren. II. Grundriss der organisation der Siphonophoren. in Zoologischen Jahrbücher. Abteilung für Anatomie und Ontogenie 1018 der Tiere Abteilung für Anatomie und Ontogenie der Tiere (ed. Spengel, J. W.) (Verlag Von Gustav, Iena, 1896).

41. Chun, C. Über die cyklische Entwickelung der Siphonophoren. Sitzungsberichte der Königlich Preussischen Akademie der Wissenschaften zu Berlin 1885, 511–529 (1885).

42. Grimmelikhuijzen, C. J. P., Spencer, A. N. & Carré, D. Organization of the nervous system of physonectid siphonophores. Cell Tissue Res 246, 463–479 (1986).

43. Siebert, S. et al. Stem cells in *Nanomia bijuga* (Siphonophora), a colonial animal with localized growth zones. Evodevo 6, 22 (2015).

44. Moser, F. Die Siphonophoren der Deutschen Südpolar Expedition, 1901-3. in Deutsche Sudpolar-Expedition 1901-1903 (ed. von Drygalski, E.) (Walter De Gruyter & Co., Berlin, 1925).

45. Siebert, S., Pugh, P. R., Haddock, S. H. D. & Dunn, C. W. Re-evaluation of characters in Apolemiidae (Siphonophora), with description of two new species from Monterey Bay, California. Zootaxa 3702, 201–232 (2013).

46. McClain, C. R. et al. Navigating uncertainty in maximum body size in marine metazoans. Ecol Evol 14, e11506 (2024).

47. Hiscock, K., Mapstone, G. M., Conway, D. V. P. & Halliday, N. Occurrence of the physonect siphonophore *Apolemia uvaria* off Plymouth and in south-west England. Mar Biodivers Rec 3, 3–6 (2010).

48. Mackie, G. O. Coordinated behavior in hydrozoan colonies. in Animal colonies. Development and Function through time (eds. Boardman, R. S., Cheetham, A. H. & Oliver Jr., W. A.) (Dowden, Hutchingon & Ross, Inc., Pennsylvania, 1973).

49. Leclère, L. & Röttinger, E. Diversity of cnidarian muscles: Function, anatomy, development and regeneration. Front Cell Dev Biol 4, 157 (2017).

50. Holz, O. et al. Bud Detachment in Hydra Requires Activation of Fibroblast Growth Factor Receptor and a Rho-ROCK-Myosin II Signaling Pathway to Ensure Formation of a Basal Constriction. Developmental Dynamics 246, 502–516.

51. Sinigaglia, C. et al. Pattern regulation in a regenerating jellyfish. Elife 9, 1–33 (2020).

52. Connally, N., Anderson, C. P., Bolton, J. E., Bolton, E. W. & Buss, L. W. The Selective Myosin II Inhibitor Blebbistatin Reversibly Eliminates Gastrovascular Flow and Stolon Tip Pulsations in the Colonial Hydroid Podocoryna carnea. PLoS One 10, e0143564 (2015).

53. Biggs, D. C. Field studies of fishing, feeding, and digestion in siphonophores. Mar Behav Physiol 4, 261–274 (1977).

54. Grossmann, M. M., Collins, A. G. & Lindsay, D. J. Description of the eudoxid stages of *Lensia havock* and *Lensia leloupi* (Cnidaria: Siphonophora: Calycophorae), with a review of all known *Lensia* eudoxid bracts. Syst Biodivers 12, 163–180 (2014).

55. Marshall, D. J. & Connallon, T. Carry-over effects and fitness trade-offs in marine life histories: The costs of complexity for adaptation. Evol Appl 16, 474–485 (2023).

56. Dunn, C. W., Pugh, P. R. & Haddock, S. H. D. Molecular phylogenetics of the siphonophora (Cnidaria), with implications for the evolution of functional specialization. Syst Biol 54, 916–935 (2005).

57. Damian-Serrano, A., Haddock, S. H. D. & Dunn, C. W. The evolution of siphonophore tentilla for specialized prey capture in the open ocean. Proceedings of the National Academy of Sciences 118, e2005063118 (2021).

58. Katoh, K., Misawa, K., Kuma, K. & Miyata, T. MAFFT: a novel method for rapid multiple sequence alignment based on fast Fourier transform. Nucleic Acids Res 15, 3059–3066 (2002).

59. Nguyen, L.-T., Schmidt, H. A., von Haeseler, A. & Minh, B. Q. IQ-TREE: A Fast and Effective Stochastic Algorithm for Estimating Maximum-Likelihood Phylogenies. Mol Biol Evol 32, 268–274 (2014).

60. Maddison, W. P. & Maddison, D. R. Mesquite: a modular system for evolutionary analysis. Version 3.81. http://www.mesquiteproject.org. (2023).

