## Supplementary materials for "The evolution of an individual-like dispersive stage in colonial siphonophores"

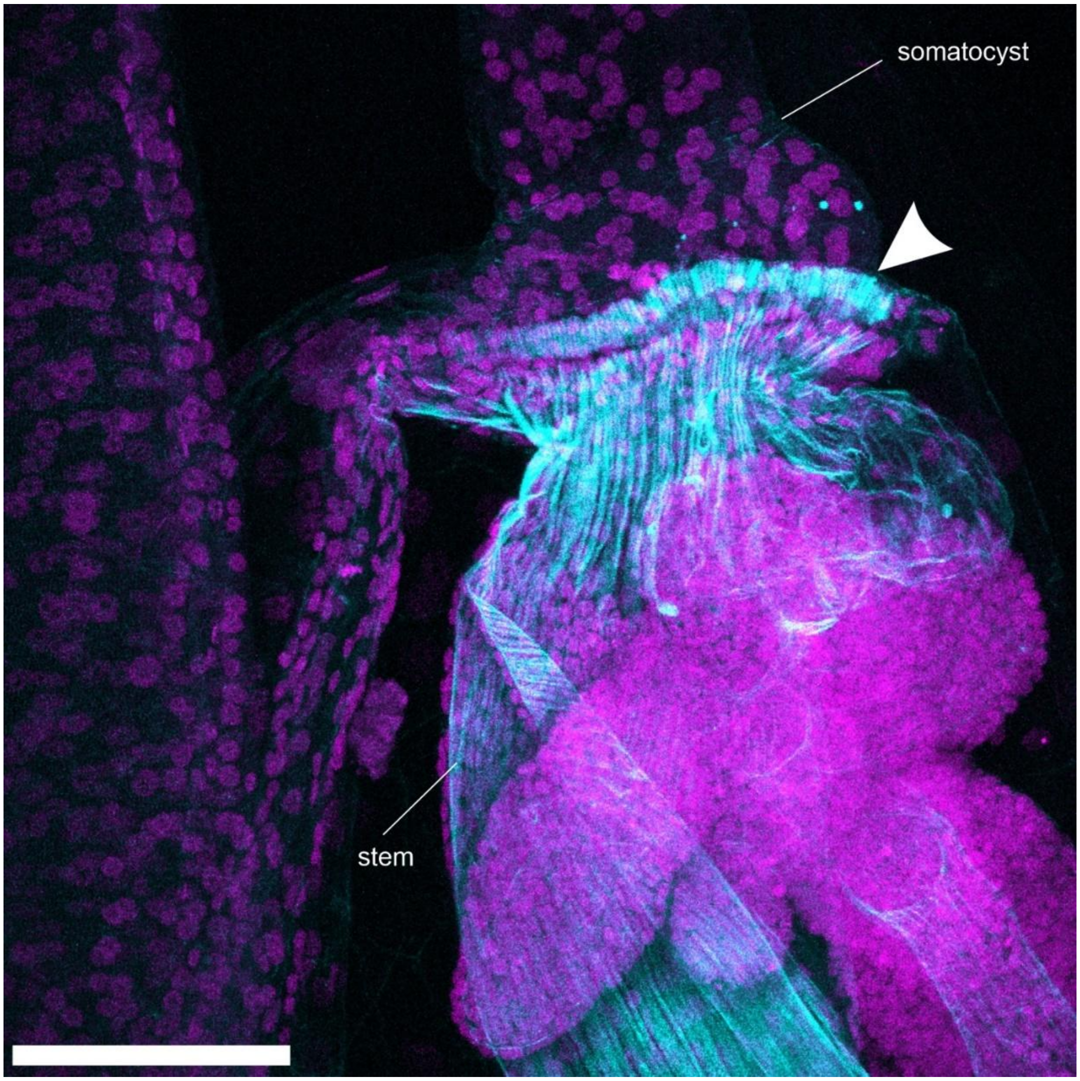


**Supplementary Fig. 1. Actin-rich structures in the stem – nectophore connection site in *Chelophyes appendiculata***

Actin (cyan) and nuclei (magenta) detection at the connection site between stem and nectophore (nectophore somatocyst indicated with label). Arrowhead point at the actin-rich structures at the connection site. Scale bar: 100 µm.

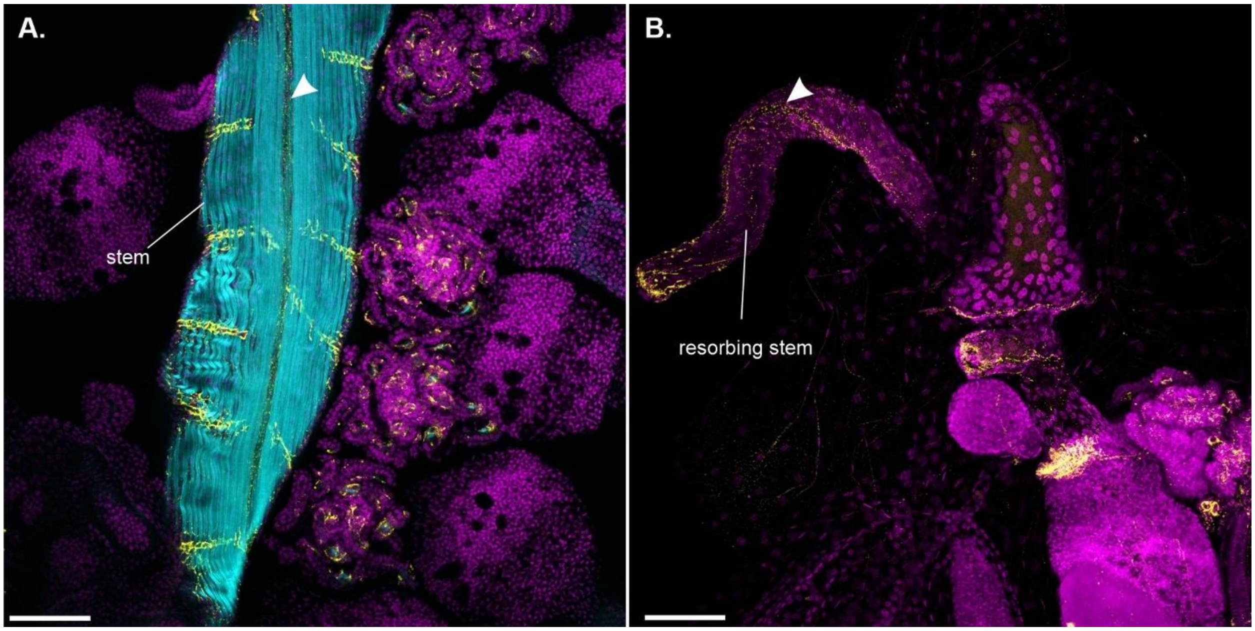


**Supplementary Fig. 2. Nervous architecture in *Chelophyes appendiculata*.**

A. Immunohistostaining of FMFRamide (yellow), actin (cyan) and nuclei (magenta) in *C. appendiculata* stem. B. Immunohistostaining of FMFRamide (yellow) and nuclei (magenta) in newly released eudoxid with resorbing stem. Arrowheads point at the stems' giant axons. Scale bars 100 µm.

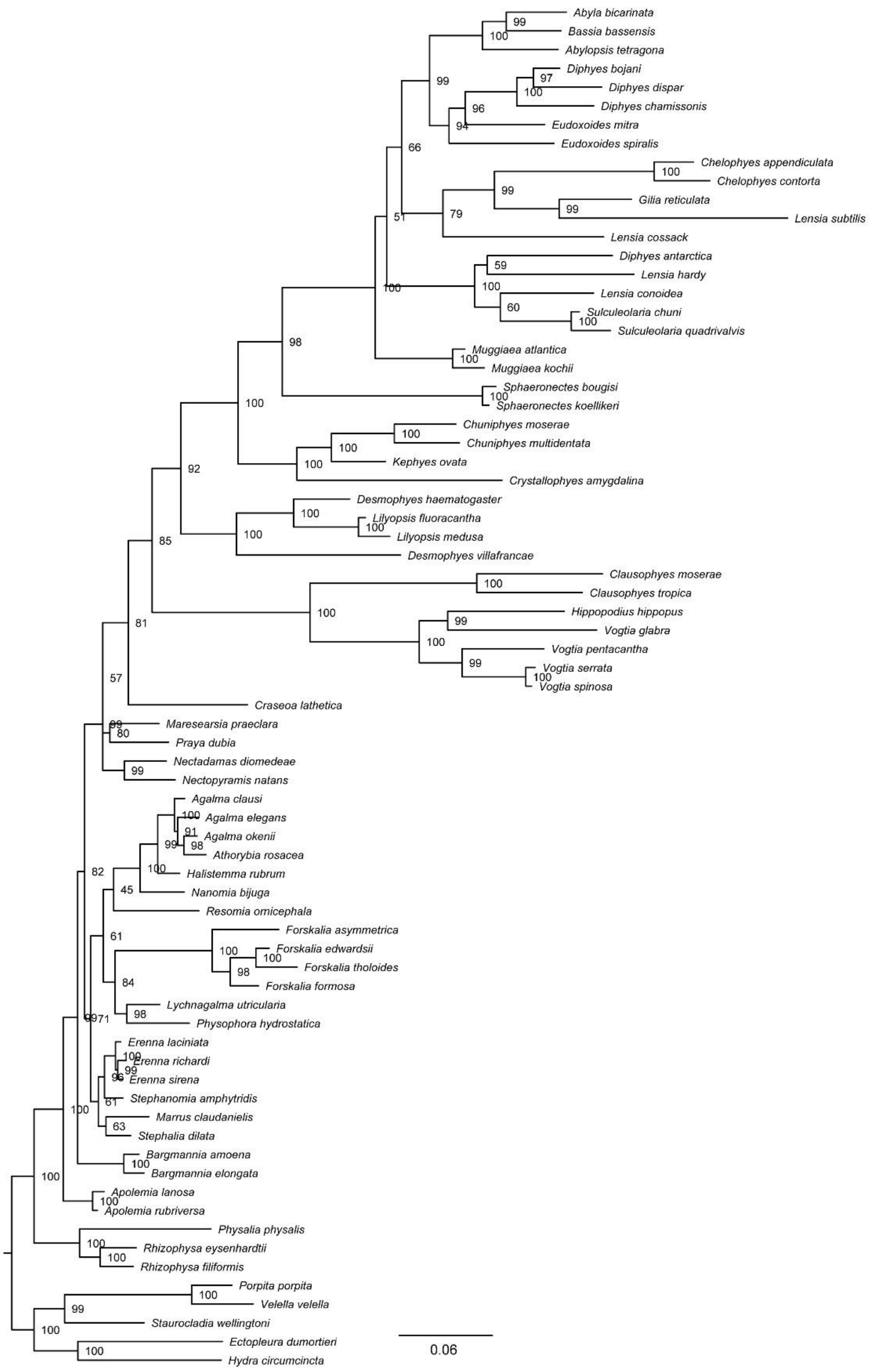


**Supplementary Fig. 3. Maximum likelihood IQTree inference, unconstrained. Node labels are bootstrap support values.**

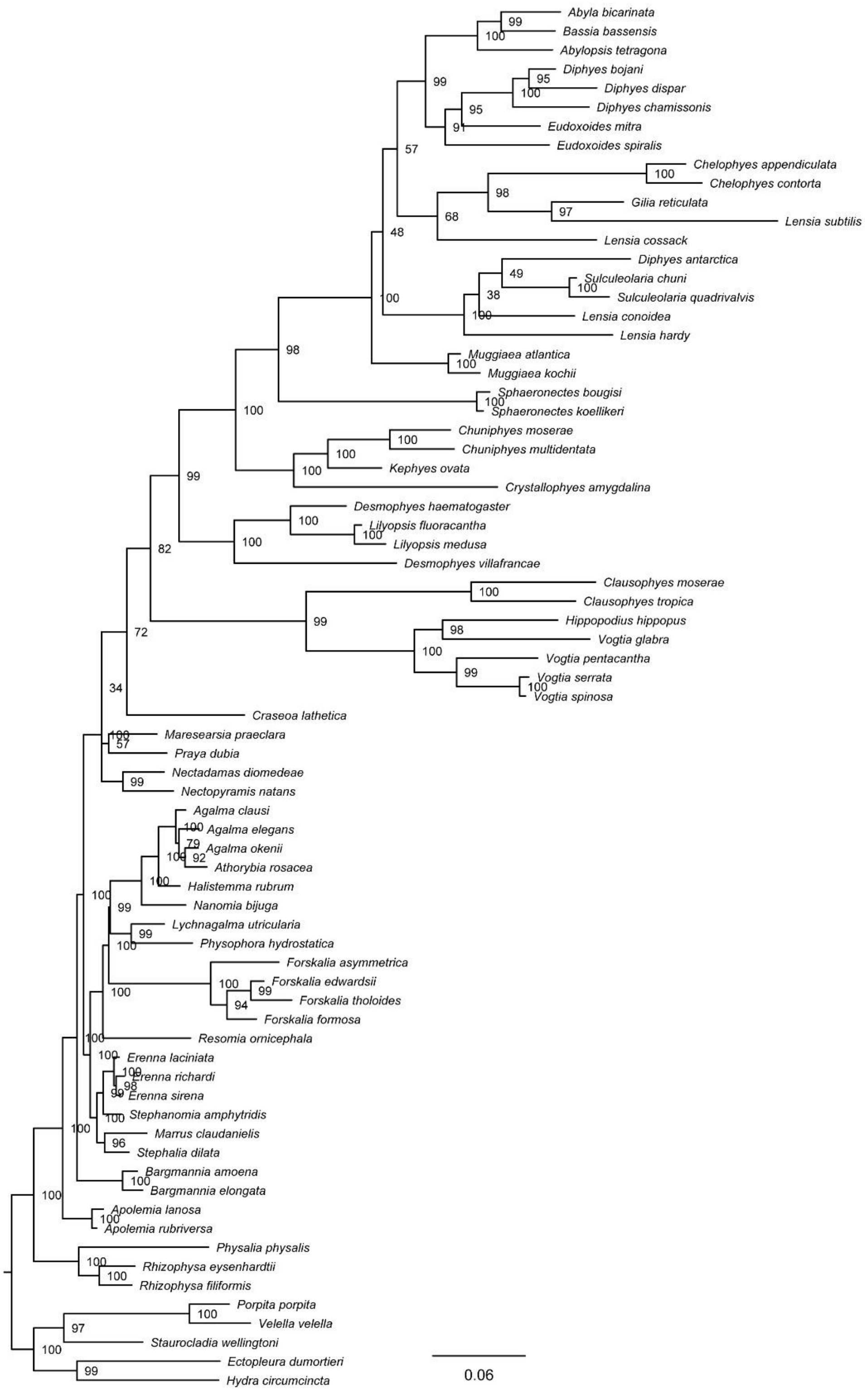


**Supplementary Fig. 4. Maximum likelihood IQTree inference, constrained. Node labels are bootstrap support values.**

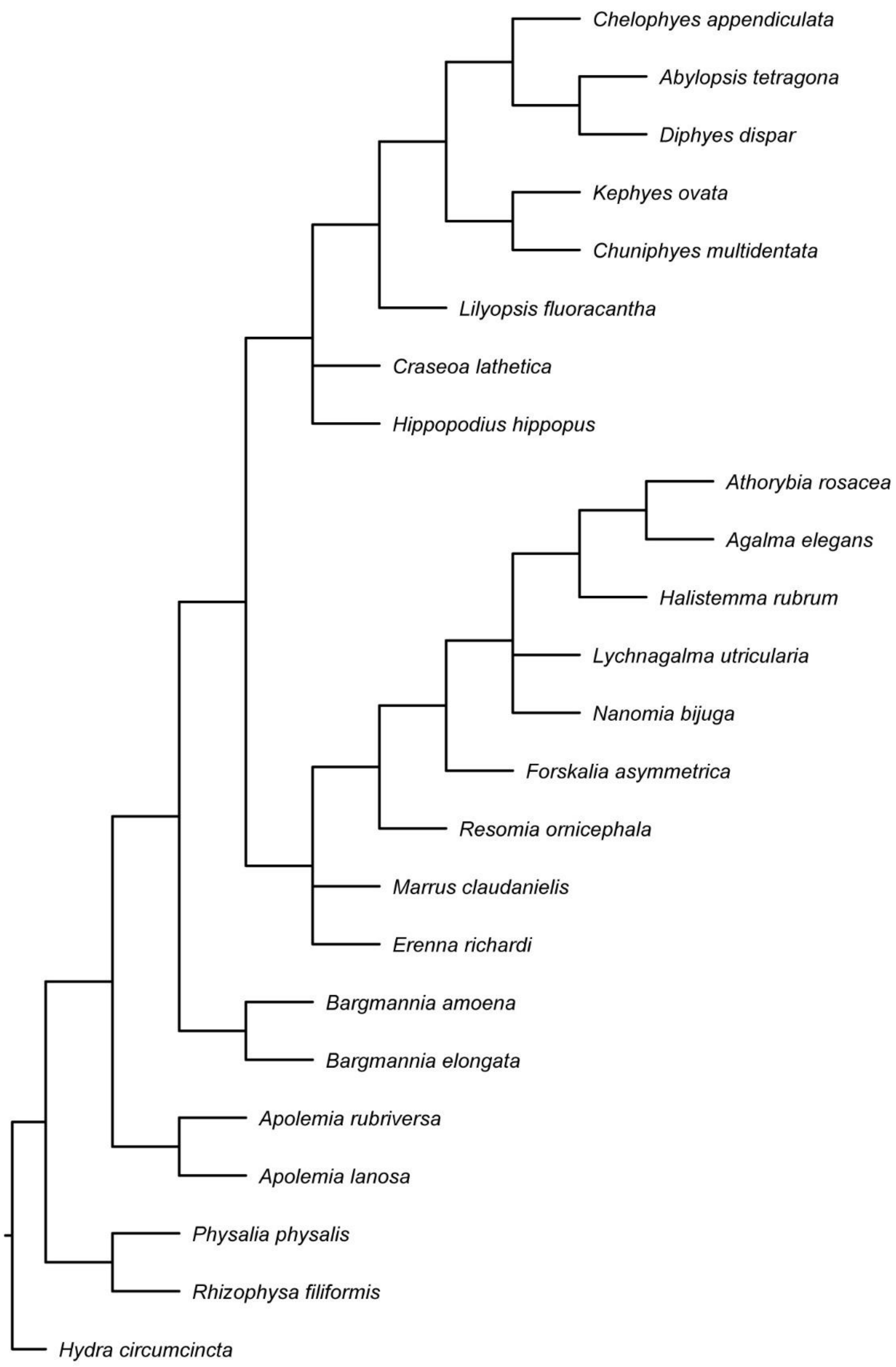


**Supplementary Fig. 5. Topology used to constrain the IQTree ML analyses from Munro et al.**[30]

**Supplementary Table 1. Observations of cormidium release from entire colonies.**

Results of observations of the cormidium release in *Chelophyes appendiculata* colonies. For details see Methods section. NA - not available.

| ID | Treatment | Stage of posterior-most cormidium | Stage of 1st cormidium released | Stage of 2nd cormidium released | Eudoxids released | Observation duration |
|---|---|---|---|---|---|---|
| 2021.3 | unperturbed | S2 | - | - | No | 48h |
| 2021.3 | unperturbed | S2 | - | - | No | 48h |
| 2021.5 | immobilized | S2 | - | - | No | 48h |
| 2021.5 | immobilized | S2 | - | - | No | 48h |
| 2021.Col2 | unperturbed | S2 | - | - | No | 24h |
| 2021.8 | immobilized | S2 | - | - | No | 24h |
| 2021.8 | immobilized | S2 | - | - | No | 24h |
| 2021.16 | immobilized | S2 | - | - | No | 24h |
| 2021.16 | immobilized | S2 | - | - | No | 24h |
| 2021.18 | unperturbed | S2 | - | - | No | 24h |
| 2021.18 | unperturbed | S2 | - | - | No | 24h |
| 2021.14.1 | immobilized | S3 | S4 | S4 | Yes | 48h |
| 2021.14.1 | immobilized | S3 | S4 | S4 | Yes | 48h |
| 2021.14.1 | immobilized | S3 | S4 | S4 | Yes | 48h |
| 2021.14.2 | immobilized | S3 | S4 | S4 | Yes | 48h |
| 2021.14.2 | immobilized | S3 | S4 | S4 | Yes | 48h |
| 2021.15 | immobilized | S3 | NA | NA | Yes | 48h |
| 2021.15 | immobilized | S3 | NA | NA | Yes | 48h |
| 2021.15 | immobilized | S3 | NA | NA | Yes | 48h |
| 2021.Col1 | unperturbed | S3 | S4 | S4 | Yes | 48h |
| 2021.6 | unperturbed | S3 | S4 | S4 | Yes | 48h |

**Supplementary Table 2. Stem fragmentation experiments.**

Data acquired from stem fragmentation experiments, listing experiment identifier (ID), colony identifier (ColonyID), type of fragments expressed as CLSF (cormidium-long-stem-fragment), staging of individual cormidia within the fragment, number of eudoxids released and observation frequency.

| ID | Colony ID | CLSF type | Cormidium staging at T0 [posterior to anterior] | Number of eudoxids | Observation frequency |
|---|---|---|---|---|---|
| 2021.18 | 2021.18.1 | 4 | S3-S3-S3-S3 | 4 | after 24h |
| 2021.18 | 2021.18.2 | 4 | S3-S3-S3-S3 | 4 | after 24h |
| 2021.19 | 2021.19.1 | 4 | S3-S3-S3-S3 | 4 | after 24h |
| 2021.19 | 2021.19.2 | 4 | S3-S3-S3-S3 | 4 | after 24h |
| 2021.19 | 2021.19.3 | 4 | S3-S3-S3-S3 | 4 | after 24h |
| 2021.19 | 2021.19.4 | 4 | S3-S3-S3-S3 | 4 | after 24h |
| 2021.19 | 2021.19.5 | 4 | S3-S3-S3-S3 | 4 | after 24h |
| 2021.2 | 2021.2.1 | 4 | S2-S2-S2-S2 | 0 | after 24h |
| 2021.2 | 2021.2.1 | 4 | S2-S2-S2-S2 | 0 | after 24h |
| 2021.2 | 2021.2.2 | 4 | S2-S2-S2-S2 | 0 | after 24h |
| 2021.2 | 2021.2.3 | 4 | S2-S2-S2-S2 | 0 | after 24h |
| 2021.2 | 2021.2.4 | 4 | S2-S2-S2-S2 | 0 | after 24h |
| 2021.2 | 2021.2.5 | 4 | S2-S2-S2-S2 | 0 | after 24h |
| 2021.2 | 2021.2.6 | 4 | S2-S2-S2-S2 | 0 | after 24h |
| 2021.2 | 2021.2.7 | 4 | S2-S2-S2-S2 | 0 | after 24h |
| 2021.3 | 2021.3.1 | 4 | S2-S2-S2-S2 | 0 | every 24h for 3 days |
| 2021.3 | 2021.3.2 | 4 | S2-S2-S2-S2 | 0 | every 24h for 3 days |
| 2021.4 | 2021.4.1 | 4 | S2-S2-S2-S2 | 0 | after 2, 8, 18, 23, 25h |
| 2021.4 | 2021.4.2 | 4 | S2-S2-S2-S2 | 0 | after 2, 8, 18, 23, 25h |
| 2022.1 | 2022.1.1 | 2 | S4-S4 | 2 | every 2h for 8h, then occasionally |
| 2022.1 | 2022.1.2 | 3 | S2-S2-S2 | 0 | every 2h for 8h, then occasionally |
| 2022.1 | 2022.1.3 | 3 | S4-S4-S4 | 3 | every 2h for 8h, then occasionally |
| 2022.1 | 2022.1.3 | 3 | S4-S4-S3 | 3 | every 2h for 8h, then occasionally |
| 2022.1 | 2022.1.4 | 3 | S4-S3-S3 | 3 | every 2h for 8h, then occasionally |
| 2022.1 | 2022.1.4 | 3 | S3-S3-S3 | 3 | every 2h for 8h, then occasionally |
| 2022.4 | 2022.4.1 | 3 | S3-S3-S3 | 3 | every 1h for 5h, then every 24h |

**Supplementary Table 3. Pharmacological inhibition experiment**

Summary of results of pharmacological inhibition experiments, experiment identifier (ID), type of fragments expressed as CLSF (cormidium-long-stem-fragment), staging of individual cormidia within the fragment and five scored phenotypes: released cormidia, remodeled bracts, aberrant bract morphology, aberrant bracteal canal, stem resorbed.

| ID | CLSF type | Treatment | Cormidium staging at T0 [posterior to anterior] | Released cormidia | Remodeled bracts | Aberrant bract morphology | Aberrant bracteal canal | Stems resorbed |
|---|---|---|---|---|---|---|---|---|
| 2022.5.inh | 3 | 1 µM B | S4-S4-S4 | 3 | 0 | 3 | 3 | 0 |
| 2022.5.inh | 3 | 1 µM B | S3-S3-S3 | 3 | 0 | 3 | 3 | 0 |
| 2022.5.inh | 3 | 1 µM B | S3-S3-S3 | 1 | 0 | 3 | 3 | 0 |
| 2022.bb.inh | 3 | 1 µM B | S4-S4-S4 | 3 | 0 | 3 | 3 | 0 |
| 2022.bb.inh | 3 | 1 µM B | S2-S2-S2 | 0 | 0 | 3 | 3 | 0 |
| 2022.fs.inh | 2 | 1 µM B | S3-S3 | 0 | 0 | 2 | 2 | 0 |
| 2022.fs.inh | 2 | 1 µM B | S2-S2 | 0 | 0 | 2 | 2 | 0 |
| 2022.5.inh | 3 | 5 µM B | S4-S4-S4 | 1 | 0 | 3 | 3 | 0 |
| 2022.5.inh | 3 | 5 µM B | S3-S3-S3 | 1 | 0 | 3 | 3 | 0 |
| 2022.5.inh | 3 | 5 µM B | S3-S3-S3 | 0 | 0 | 3 | 3 | 0 |
| 2022.bb.inh | 3 | 5 µM B | S4-S4-S4 | 3 | 0 | 3 | 3 | 0 |
| 2022.bb.inh | 3 | 5 µM B | S3-S2-S2 | 1 | 0 | 3 | 3 | 0 |
| 2022.fs.inh | 2 | 5 µM B | S3-S3 | 0 | 0 | 2 | 2 | 0 |
| 2022.fs.inh | 2 | 5 µM B | S2-S2 | 0 | 0 | 2 | 2 | 0 |
| 2022.5.inh | 3 | DMSO | S4-S4-S4 | 3 | 3 | 0 | 0 | 3 |
| 2022.5.inh | 3 | DMSO | S3-S3-S3 | 3 | 3 | 0 | 0 | 3 |
| 2022.fs.inh | 2 | DMSO | S3-S3 | 2 | 2 | 0 | 0 | 2 |
| 2022.fs.inh | 2 | DMSO | S2-S2 | 0 | 0 | 0 | 0 | 0 |
| 2022.5.inh | 3 | MFSW | S4-S4-S4 | 3 | 3 | 0 | 0 | 3 |
| 2022.5.inh | 3 | MFSW | S3-S3-S3 | 3 | 3 | 0 | 0 | 3 |
| 2022.bb.inh | 3 | MFSW | S4-S4-S4 | 3 | 3 | 0 | 0 | 3 |
| 2022.bb.inh | 3 | MFSW | S2-S2-S2 | 0 | 0 | 0 | 0 | 0 |
| 2022.fs.inh | 2 | MFSW | S3-S3 | 2 | 2 | 0 | 0 | 2 |
| 2022.fs.inh | 2 | MFSW | S2-S2 | 0 | 0 | 0 | 0 | 0 |
| 2022.2.inh | 2 | $MgCl_2$ | S3-S3 | 0 | 2 | 0 | 0 | 0 |

**Supplementary Table 4. List of sequences used in this study, with their GenBank accession numbers. Novel sequences generated here are written in bold.**

| Sub-order | Family | Species | 16S accession number | 18S accession number |
|---|---|---|---|---|
| Calycophorae | Abylidae | *Abyla bicarinata* | **PV190272** | **PV138272** |
| Calycophorae | Abylidae | *Abylopsis tetragona* | AY935303 | AY937345.1 |
| Calycophorae | Abylidae | *Bassia bassensis* | **PV190273** | **PV138273** |
| Calycophorae | Clausophyidae | *Chuniphyes moserae* | **PV190276** | **PV138276** |
| Calycophorae | Clausophyidae | *Chuniphyes multidentata* | AY935293 | AY937335.1 |
| Calycophorae | Clausophyidae | *Clausophyes moserae* | **PV190277** | **PV138277** |
| Calycophorae | Clausophyidae | *Clausophyes tropica* | **PV190278** | **PV138278** |
| Calycophorae | Clausophyidae | *Crystallophyes amygdalina* | KX374466 | KX421850.1 |
| Calycophorae | Clausophyidae | *Kephyes ovata* | AY935294 | AY937336.1 |
| Calycophorae | Diphyidae | *Chelophyes appendiculata* | **PV190274** | **PV138274** |
| Calycophorae | Diphyidae | *Chelophyes contorta* | **PV190275** | **PV138275** |
| Calycophorae | Diphyidae | *Diphyes antarctica* | **PV190280** | **PV138280** |
| Calycophorae | Diphyidae | *Diphyes bojani* | MZ230468.1 | MZ230443.1 |
| Calycophorae | Diphyidae | *Diphyes chamissonis* | MZ230470.1 | MZ230445.1 |
| Calycophorae | Diphyidae | *Diphyes dispar* | AY935276 | AY937318.1 |
| Calycophorae | Diphyidae | *Eudoxoides mitra* | MZ230478.1 | MZ230453.1 |
| Calycophorae | Diphyidae | *Eudoxoides spiralis* | **PV190281** | **PV138281** |
| Calycophorae | Diphyidae | *Gilia reticulata* | KX374469.1 | KX421854.1 |
| Calycophorae | Diphyidae | *Lensia conoidea* | AY935318 | AY937360.1 |
| Calycophorae | Diphyidae | *Lensia cossack* | MZ230483.1 | MZ230458.1 |
| Calycophorae | Diphyidae | *Lensia hardy* | **PV190282** | **PV138282** |
| Calycophorae | Diphyidae | *Lensia subtilis* | **PV190283** | **PV138273** |
| Calycophorae | Diphyidae | *Muggiaea atlantica* | AY935295 | AY937337.1 |
| Calycophorae | Diphyidae | *Muggiaea kochii* | **PV190285** | **PV138286** |
| Calycophorae | Diphyidae | *Sulculeolaria chuni* | **PV190287** | **PV138288** |
| Calycophorae | Diphyidae | *Sulculeolaria quadrivalvis* | **PV190288** | **PV138289** |
| Calycophorae | Hippopodiidae | *Hippopodius hippopus* | AY935299 | AF358069.1 |
| Calycophorae | Hippopodiidae | *Vogtia glabra* | AY935308 | AY937350.1 |
| Calycophorae | Hippopodiidae | *Vogtia pentacantha* | AY935320 | AY937362.1 |
| Calycophorae | Hippopodiidae | *Vogtia serrata* | **PV190289** | **PV138290** |
| Calycophorae | Hippopodiidae | *Vogtia spinosa* | **PV190290** | **PV138291** |
| Calycophorae | Prayidae | *Craseoa lathetica* | AY935297 | AY937339.1 |
| Calycophorae | Prayidae | *Desmophyes haematogaster* | DQ080006 | KX421851.1 |
| Calycophorae | Prayidae | *Desmophyes villafrancae* | **PV190279** | **PV138279** |
| Calycophorae | Prayidae | *Lilyopsis fluoracantha* | SRR1548373 | AY919607.1 |
| Calycophorae | Prayidae | *Lilyopsis medusa* | | **PV138284** |
| Calycophorae | Prayidae | *Maresearsia praeclara* | **PV190284** | **PV138285** |
| Calycophorae | Prayidae | *Nectadamas diomedeae* | AY935306 | AY937348.1 |

| | | | | |
|---|---|---|---|---|
| Calycophorae | Prayidae | *Nectopyramis natans* | AY935307 | AY937349.1 |
| Calycophorae | Prayidae | *Praya dubia* | AY935285 | AY937326.1 |
| Calycophorae | Sphaeronectidae | *Sphaeronectes bougisi* | **PV190286** | **PV138287** |
| Calycophorae | Sphaeronectidae | *Sphaeronectes koellikeri* | AY935301 | AF358070.1 |
| Cystonectae | Physaliidae | *Physalia physalis* | AY935284 | AF358065.1 |
| Cystonectae | Rhizophysidae | *Rhizophysa eysenhardtii* | AY935309 | AY937351.1 |
| Cystonectae | Rhizophysidae | *Rhizophysa filiformis* | AY935286 | AY937327.1 |
| Physonectae | Agalmatidae | *Agalma clausi* | AY935270 | AY937312.1 |
| Physonectae | Agalmatidae | *Agalma elegans* | AY935271 | AY937313.1 |
| Physonectae | Agalmatidae | *Agalma okenii* | AY935272 | AY937314.1 |
| Physonectae | Agalmatidae | *Athorybia rosacea* | AY935274 | AY937316.1 |
| Physonectae | Agalmatidae | *Halistemma rubrum* | AY935316 | AY937323.1 |
| Physonectae | Agalmatidae | *Lychnagalma utricularia* | DQ080009 | SRR1548374 |
| Physonectae | Agalmatidae | *Marrus claudanielis* | DQ080007 | SRR1548375 |
| Physonectae | Agalmatidae | *Nanomia bijuga* | AY935296 | AY937338.1 |
| Physonectae | Apolemiidae | *Apolemia lanosa* | KF214712 | KF214714.1 |
| Physonectae | Apolemiidae | *Apolemia rubriversa* | KF214713 | KF214715.1 |
| Physonectae | Erennidae | *Erenna laciniata* | KX752722 | KX752704.1 |
| Physonectae | Erennidae | *Erenna richardi* | KX752723 | KX752705.1 |
| Physonectae | Erennidae | *Erenna sirena* | KX752725 | AY937361.1 |
| Physonectae | Forskaliidae | *Forskalia asymmetrica* | AY935277 | AY937319.1 |
| Physonectae | Forskaliidae | *Forskalia edwardsii* | AY935278 | AY937320.1 |
| Physonectae | Forskaliidae | *Forskalia formosa* | AY935302 | AY937344.1 |
| Physonectae | Forskaliidae | *Forskalia tholoides* | AY935279 | AY937321.1 |
| Physonectae | Physophoridae | *Physophora hydrostatica* | AY935300 | AY937342.1 |
| Physonectae | Pyrostephidae | *Bargmannia amoena* | AY935292 | AY937333.1 |
| Physonectae | Pyrostephidae | *Bargmannia elongata* | AY935321 | AY937334.1 |
| Physonectae | Resomiidae | *Resomia ornicephala* | SRR1548382 | SRR1548382 |
| Physonectae | Rhodaliidae | *Stephalia dilata* | AY935315 | AY937357.1 |
| Physonectae | Stephanomidae | *Stephanomia amphytridis* | AY935280 | AY937322.1 |
| Outgroup | | *Ectopleura dumortieri* | EU305474 | EU272616.1 |
| Outgroup | | *Hydra circumcincta* | GU722773 | EU876568.1 |
| Outgroup | | *Porpita porpita* | AY935322 | AF358086.1 |
| Outgroup | | *Staurocladia wellingtoni* | AJ580934 | AF358084.1 |
| Outgroup | | *Velella velella* | AY935323 | AF358087.1 |

**Supplementary Table 5. Summary of buoyancy experiments**

Data were summarized to show the number of cormidia within each colony tested (ColonyID) and the outcome of experiment as percent of cormidia at a given position (bottom or floating) at the begging of experiment (T = 0h) and after 24h

| Colony ID | Number of cormidia number | Position at T = 0h | | Position at T = 24h | |
|---|---|---|---|---|---|
| | | % at the bottom | % floating | % at the bottom | % floating |
| col.1 | 25 | 100 | 0 | 0 | 100 |
| col.2 | 27 | 100 | 0 | 0 | 100 |
| col.3 | 13 | 100 | 0 | 0 | 100 |
| col.4 | 14 | 100 | 0 | 0 | 100 |
| col.5 | 10 | 100 | 0 | 0 | 100 |

**Supplementary Data 1. Character matrix used in character state evolution reconstruction.**

**Supplementary Data 2. Literature review on ecological mismatch between eudoxids and colonies with full reference list.**
Dataset contains 1) results of literature review, indicating data type per reference analyzed, observation on mismatch and species to which this mismatch pertains, 2) full references to 1), 3) information on data used to generate Fig. 8B-E., 4-7) Data used to generate Fig. 8B-E.

**Supplementary Data 3. Nexus file for the morphological evolution analysis with Mesquit**

**Supplementary Data 4. Concatenated alignment of 16S and 18S sequences**

**Supplementary Data 5. Code used to generate plots in Fig. 6 and 8.**

**Supplementary Movie 1. Video documentation of eudoxid release**.
Movie showing detachment ring constriction in 3-cormidia long stem fragment of *Chelophyes appendiculata* and subsequent detachment of cormidia, followed by eudoxid maturation.

**Supplementary Movie 2. Video documentation of eudoxid release.**
Movie showing sequential fragmentation and eudoxid maturation originating from 3-cormidia long stem fragment of *Chelophyes appendiculata*.

**Supplementary Movie 3. Video documentation of eudoxid maturation.**
Movie showing eudoxid maturation processes that the single cormidium undergoes following its separation from the colony.

**Supplementary Movie 4. Video documentation of veronica movement in *Chelophyes appendiculata* eudoxid.**

**Supplementary Movie 5. Video documentation of veronica movement in *Chelophyes appendiculata* eudoxid.**
